# Loss of ELM1B impairs mitochondrial fission, matrix redox state and stress tolerance in *Physcomitrium patens*

**DOI:** 10.64898/2026.09.01.748533

**Authors:** Sadia S. Tamanna, Sophie Pompejus, Subasini Thangamani, Cloé Gadoud, Isabelle Nermerich, Timo Mühlhaus, Stefanie J. Müller-Schüssele

## Abstract

Mitochondria are endosymbiont-derived organelles that play a central role in cellular metabolism, energy production and stress responses. While single mitochondria represent functional units, they continuously exchange their contents through fusion and fission, facing stress conditions as a dynamic population. To date, it remains largely unknown how stress alters mitochondrial dynamics in plants and how altered dynamics affect mitochondrial properties and plant stress resilience.

Here, we investigate mitochondrial dynamics in response to oxidative stress in the non-vascular model plant *Physcomitrium patens*. By creating mutants with impaired mitochondrial fission in different reporter lines for mitochondrial parameters, we additionally analyse effects of chronic changes to mitochondrial population dynamics. We found that Mito-Paraquat (MtPQ) treatment increased the glutathione redox potential *E*_GSH_ in mitochondria, the cytosol and chloroplasts, as monitored via roGFP2-based genetically encoded biosensors. Mitochondria elongated within hours and showed a concomitant and heterogenous increase of matrix EOS_red_, that we propose as a marker for matrix protein damage. Mitochondrial fission mutants lacking PpELM1B (ELONGATED MITOCHONDRIA) displayed distinct changes of mitochondrial morphology parameters as determined by automated 3D-segmentation and feature mapping (MorphoMapper) of confocal z-stacks. Elongated mitochondria in *Ppelm1b^ge^*lines showed an oxidative matrix *E*_GSH_ shift and increased matrix EOS_red_ while matrix mixing still occurred, albeit at the same slow rate as in wildtype, within days. Macroscopically, *Ppelm1b^ge^* lines displayed reduced growth, decreased respiration, and a higher sensitivity to oxidative stress.

Our results show that plant mitochondrial morphology and physiological parameters specifically shift in response to stress and impaired fission. Mitochondrial fission is vital to maintain a healthy mitochondrial population that sustains plant oxidative stress tolerance.

## INTRODUCTION

Mitochondria as endosymbiont-derived organelles are vital components of eukaryotic life (Archibald, 2015; Dyall *et al*., 2004). Mitochondrial processes drive energy metabolism, are involved in disease, defense, aging and plant senescence as well as contribute to essential biosynthesis pathways (Van Aken, 2021; Liesa and Shirihai, 2013; Suomalainen and Nunnari, 2024). Mitochondria share common features with the second endosymbiont-derived organelle, plastids, such as organellar DNA and propagation by division. In contrast, only mitochondria show fusion events with bulk transfer and mixing of matrix content and even cristae (Arimura, 2018; Schattat *et al*., 2012; Wilkens *et al*., 2013). The structure and dynamics of a cell’s population of mitochondria is increasingly characterized and perceived as a response factor in metabolic adjustments, connectivity to other subcellular compartments as well as stress responses (A., H., Khan *et al*., 2024; Ryu *et al*., 2024; Wikstrom *et al*., 2009). Thus, mitochondrial population dynamics respond to altered metabolic demands, cellular specialization/differentiation, as well as can suffer from or create heterogeneity within cellular (sub)populations. This comprises the existence and distribution of different mtDNA molecules (heteroplasmy) (Aryaman *et al*., 2019; Giannakis *et al*., 2022), and extents to physiological properties and protein content (Ngo *et al*., 2021; Vazquez-Calvo *et al*., 2023) with likely many yet unknown factors influencing mitochondrial heterogeneity. While first local responses occur at the single organelle level, many factors regulate and influence the ensuing population level response (Taskin *et al*., 2026; Liesa and Shirihai, 2013; Casler and Lackner, 2025). With the emergence of AI-assisted and computationally driven automated image analysis, the characterization and analysis of mitochondrial morphology on a population scale have become feasible (Chustecki *et al*., 2021; Ahmad *et al*., 2013; Lefebvre *et al*., 2021; Thangamani *et al*., 2026).

Plant mitochondria exist as a ‘discontinuous whole’ with several hundred spherical to slightly elongated physically discrete organelles (Logan, 2006a; Logan, 2006b; Chustecki *et al*., 2021; Furt *et al*., 2012), while more network-like fused states are described for dividing and meristematic cells (Sheahan *et al*., 2005; Seguí-Simarro *et al*., 2008). In contrast to animal and fungal cells, many aspects of plant mitochondria population biology remain unknown and have been mainly studied in few flowering plant model species.

Using the model moss *Physcomitirum patens*, we address how plant mitochondrial populations respond to oxidative stress and whether fission dynamics are important for matrix exchange and mixing at a population level. We sense and dynamically follow physiological parameters and protein damage to link observed population behavior to functional aspects.

## MATERIAL AND METHODS

### Plant material and growth conditions

*Physcomitrium patens* (Medina *et al*., 2019) (*Physcomitrella patens* (Hedw.) Bruch & Schimp. ecotype Gransden 2004; International Moss Stock Center (IMSC, http://www.moss-stock-center.org), accession number 40001) was used as the wildtype (WT) background. Moss culture was maintained axenically in liquid KNOP ME medium or on KNOP ME agar plates. KNOP medium contained KH_2_PO_4_ (250 mgL^-1^), KCl (250 mgL^-1^), MgSO_4_ x 7H_2_O (250 mgL^-1^), Ca(NO)_2_ x 4H_2_O (1 gL^-1^) and FeSO_4_ x 7H_2_O (12.5 mgL^-1^) supplemented with microelements (ME) containing H_3_BO_3_, MnSO_4_, ZnSO_4_, KI, Na_2_MoO_4_ x 2H_2_O, Co(NO_3_)_2_, CuSO_4_ (Reski and Abel, 1985; Egener *et al*., 2002) and was adjusted to pH 5.8. Solid medium contained 12 gL^-1^ purified agar (Oxoid). Cultures were grown at 22°C under long-day conditions (16 h light/8 h dark) at approximately 100 µmol photons m^-2^s^-1^ in a plant growth cabinet (Fitotron ® SGC 2, Weiss Technik). Protonema cultures were maintained in 500 ml Erlenmeyer flasks on a rotary shaker at approximately 100 rpm and moss cultures were regularly homogenized in fresh medium using an Ultra-Turrax T25 dispersion tool (IKA-Werke GmbH) at 14,000 rpm for 1-1.5 min. For oxidative stress treatments, 5 ml of three-day-old protonema cultures were incubated either without treatment or with 50 µM Mito-Paraquat (MtPQ, MedChem Express HY-130278) under standard growth conditions.

For growth phenotyping of genetically modified lines, colony growth was started with a single gametophore tip inserted into KNOP ME agar plates, placing gene-edited lines, corresponding background lines, and WT controls on the same plate to allow growth under identical conditions. Plates were incubated for four weeks under standard growth conditions. Colony growth was documented using a stereomicroscope (Leica M205 FCA equipped with a Leica K3C camera) with identical magnification, zoom, and imaging settings for all samples. Colony area was quantified from plate images using Fiji/ImageJ by thresholding the images and measuring colony area with the Analyze Particles function (Schindelin *et al*., 2012).

### Generation of stable transgenic and mutant lines

The mitochondrial reporter line mtmEOS #44 is described in Mueller and Reski (2015). MPP_TP_-roGFP2-hGrx1 #47 and hGrx1-roGFP2 #39 were generated in the ecotype Gransden WT background. To this end, an expression cassette composed of the maize ubiquitin 1 promoter, the respective roGFP2-based sensor coding sequence (Grx1-roGFP2 (Gutscher *et al*., 2008) and roGFP2-Grx1 (Albrecht *et al*., 2014)) and the *Agrobacterium tumefaciens* Nopalin synthase terminator was cloned using classical restriction (BamH1) between homologous regions targeting the ‘*P. patens* Targeting Locus 1 (PTA1 locus Pp1s310_4)’ (Kubo *et al*., 2013) in a pjet1.2 vector backbone. BspQ1 restriction sites were added for vector digestion to expose homologous ends for homologous recombination. The cut homologous recombination construct with the expression cassette was co-transformed with a plasmid containing the *nptII* resistance cassette (pBsNNNEV) for selection of transformed protoplasts on geneticin G418 (12.5 µg/mL) for four weeks.

Mutant lines were generated using a CRISPR/Cas9 system for *P. patens* (Mallett *et al*., 2019) in the respective reporter line backgrounds. For each *ELM1* gene, two protospacers targeting exons two and six of *PpELM1B* (*Pp3c20_13230V3.1*) and *PpELM1A* (*Pp3c23_3410V3.1*) were designed **(Table S1)**. Complete CRISPR/Cas9 expression plasmids containing the sgRNA and Cas9 expression cassettes (pMH (sgRNAs *ELM1B*), pMK (sgRNAs *ELM1A*)) were introduced into *P. patens* protoplasts by PEG-mediated transformation (Hohe *et al*., 2004). After 2-3 weeks on hygromycin B selection (12.5 µg/mL) or geneticin G418 (12.5 µg/mL), candidate plants were screened by PCR using primer pairs flanking both sgRNA target regions **(Table S1)**. Genomic DNA was extracted from gametophores according to Edwards *et al*. (1991) in buffer containing 200 mM Tris-HCl, pH 7.5, 250 mM NaCl, 25 mM EDTA and 0.5% SDS. Tissue was flash-frozen in liquid nitrogen and pulverized with glass beads for 90 s at 30 Hz using a TissueLyser II (Qiagen). When required, DNA was further purified using the NucleoSpin gDNA Clean-up kit (Macherey-Nagel). PCR was performed using Phire Hot Start II DNA Polymerase (Thermo Fisher Scientific) and analysed on 1–2% agarose gels in 1xTAE buffer containing HDGreen Plus and visualized using an INTAS ECL ChemoStar imaging system. Sanger sequencing was used to confirm mutations. Lines with frameshifts and premature stop codons were selected for further analysis.

### Confocal laser scanning microscopy and mtmEOS photoconversion

Mitochondrial morphology was analysed using protonema filaments of the mtmEOS #44 background line and corresponding *Ppelm1b^ge^* lines grown in liquid medium. In each filament, the fourth or fifth cell from the protonema tip was selected for imaging in all mtmEOS-based analyses. For repeated imaging of the same cells, protonema tissue was immobilized in 0.6% low-melting point agar prepared in KNOP ME medium in gridded µ-Slide 8 Well chamber slides (Ibidi). During imaging, samples were covered with fresh KNOP ME medium to prevent drying. Images were acquired using a Zeiss LSM 880 confocal laser scanning microscope equipped with an Axio Observer and a C-Apochromat 40×/1.2W objective with AutoCorr.. Imaging settings were chosen to balance spatial resolution with the acquisition time required for full-cell z-stacks. The zoom factor was adjusted according to individual cell size and ranged from 2.5 to 3.6. Images were acquired at 512 × 512 pixels with an averaging of 2. For each cell, the z-range and number of optical sections were determined using the Optimal setting in the ZEISS ZEN software. The green and red mtmEOS signals were collected using the same split lambda detector, so that detector settings were applied equally to both channels. MtmEOS_green_ and chlorophyll autofluorescence were excited with the 488 nm argon laser at 0.8% laser power. MtmEOS_red_ was excited with a 543 nm HeNe laser at 4.0% laser power. Emission was detected at 500–544 nm for mEOS_green_, 580–624 nm for mEOS_red_ and 657–687 nm for chlorophyll autofluorescence. Transmitted-light images were acquired simultaneously using 488 nm excitation and the PMT-T1 transmitted light detector.

For local mtmEOS photoconversion, a defined region of interest corresponding to approximately 10% of the selected cell area was irradiated with the 405 nm laser to convert green mtmEOS to the red fluorescent form. Photoconversion was done with 405 nm laser light at 50% power and 500 iterations. Before and after photoconversion, the same cell was imaged using identical acquisition settings. For quantification of photoconverted mtmEOS signal spread, ROI-based fluorescence intensity analysis was performed in Icy (De Chaumont *et al*., 2012) with the ‘ROI intensity evolution’ plugin. ROI 1 was placed over the initially photoconverted region, whereas ROI 2 covered the remaining non-photoconverted region of the same cell. Red and green mtmEOS fluorescence intensities were measured in both ROIs before and after photoconversion, and the *I*_red_/*_I_*_green_ ratio was calculated. Signal spread was assessed from the change in red-to-green ratio after photoconversion. The I_red_/I_green_ ratio was calculated for ROI 1 and ROI 2 before and after photoconversion. The change in the ratio for each ROI was determined by subtracting the value before photoconversion from the value after photoconversion. The relative spread index was calculated by dividing the change in the red-to-green ratio in ROI 2 by the corresponding change in ROI 1.

### MorphoMapper analyses

Morphometric analysis of mitochondrial populations from control and MtPQ-treated samples was done using the MorphoMapper workflow (Thangamani *et al*., 2026), which comprised three-dimensional instance segmentation followed by morphological feature quantification, feature contribution analysis and population level statistical analysis. Briefly, three-dimensional instance segmentation of the confocal image stacks was performed using a custom omnipose model implemented in this workflow, to identify individual mitochondrial objects. Morphological features were subsequently quantified for the segmented mitochondria across the dataset. In addition to these morphometric descriptors, red to green mEOS fluorescence intensity ratio (mEOS_red_/mEOS_green_) was included in the feature space. To identify which of the features contributed most to variation among mitochondrial populations, we quantified relative contribution of each feature to the differences observed between conditions.

FAIR (Findable, Accessible, Interoperable, and Reusable) principles will be adopted by organizing experimental data, analysis scripts, results and associated metadata in an Annotated Research Context (ARC) utilizing the ISA (Investigation-Study-Assay) model upon publication. All the data and scripts used for analysis will be made available in the PLANTdataHUB (Weil *et al*., 2023).

### Plate reader-based analysis of roGFP2

Oxidation of roGFP2-based redox sensors was monitored in protonema cultures using a CLARIOstar® Plus (BMG Labtech) plate reader. After dispersion, three-day-old *P. patens* protonema cultures were transferred to 96-well microplates containing KNOP ME medium. Each well contained 360 µL protonema culture. After recording five baseline cycles, the measurement was paused and 40 µL of treatment solution or KNOP ME medium was added, giving a final volume of 400 µL per well. For *in vivo* calibration, samples were treated with MtPQ, 10 mM DTT or 10 mM H_2_O_2_ to obtain fully reduced and fully oxidized sensor states, respectively. Fluorescence was measured with bottom optics in multichromatic fluorescence mode. roGFP2 fluorescence was excited at 400-10 nm and 482-16 nm, and emission was detected at 530-40 nm using an LP 504 dichroic filter. The focal height was determined by autofocus using a representative well and was set to 3.2 mm. For each measurement point, the roGFP2 excitation ratio was calculated by dividing the fluorescence intensity after excitation at 400-10 nm by the intensity after excitation at 482-16 nm. Where indicated, the degree of sensor oxidation (OxD) was calculated from the fully reduced and fully oxidized states obtained after DTT and H_2_O_2_ treatment (Meyer *et al*., 2007; Schwarzländer *et al*., 2008).

### Oxygen measurements

Oxygen concentration was measured in protonema cultures using a Clark-type Unisense OX-

200 oxygen sensor connected to a UniAmp multichannel amplifier (Unisense). For each measurement, 100 mL culture was kept under gentle stirring (Pedersen *et al*., 2013) for 2 h in darkness. Oxygen concentration was then measured during a dark-light-dark cycle of 20 min darkness, 60 min illumination and 40 min darkness. Light intensity during the light phase was 100-115 µmol photons m^-2^s^-1^. The oxygen sensor was calibrated before measurements using a zero-oxygen solution (0.1 M sodium ascorbate in 0.1 M NaOH) and air-saturated medium. Fresh weight was determined after each measurement and used to confirm comparable tissue amounts between samples.

### Statistical analysis

Statistical analyses were performed using GraphPad Prism (version 11.0.2). Ratiometric values were log-transformed before statistical analysis.

## RESULTS

### MtPQ causes oxidative stress and affects mitochondrial morphology in *P. patens*

To map the response of mitochondrial morphology to oxidative stress, we used the redox-cycler mito-paraquat (MtPQ), that mediates superoxide formation on the matrix side by re-routing electrons from respiratory complex I to molecular oxygen (Robb *et al*., 2015; K., Khan *et al*., 2024). To this end, we established treatments of protonemata grown in liquid medium, using a transgenic line expressing mitochondria-targeted photoconvertible mEOS **(Fig. 1A)**, that allows to visualise mitochondria and track matrix mixing by mitochondrial dynamics after photoconverting a fraction of the mitochondrial population (Mueller and Reski, 2015; Winter *et al*., 2026; Wiedenmann *et al*., 2004). We examined mitochondrial morphology in response to MtPQ treatment using confocal laser scanning microscopy (CLSM), generating confocal z-stacks of whole protonema cells. In control cells, mitochondria appeared mostly short and discrete, whereas MtPQ-treated cells showed visibly elongated mitochondria after 2-3 h of 50 µM MtPQ **(Fig. 1A)**. We conducted quantitative mitochondrial morphology analysis using the novel automated 3D-segmentation and feature mapping ‘MorphoMapper’ tool (Thangamani *et al*., 2026). Here, we assigned the contributions of 14 morphological features as well as the mitochondrial mEOS_red_/mEOS_green_ ratio, that allow to distinguish control and oxidative stress treatment conditions **(Fig. 1B)**. The parameter ‘skeleton length’ contributed second highest, confirming mitochondrial elongation. Notably, integrating the mitochondrial mEOS_red_/mEOS_green_ ratio as an additional parameter of the confocal microscopy data set, we found the highest contribution, overlaying the morphological changes **(Fig. 1B)**. Plotting the distribution of mEOS_red_/mEOS_green_ ratio for both control and MtPQ stress conditions, we found a broader distribution for the mitochondria populations experiencing oxidative stress, indicating heterogeneity of mitochondria within and between cells. A linear mixed-effects model accounting for image-to-image variability confirmed a significant increase in the mEOS_red_/mEOS_green_ ratio of the MtPQ-treated samples (β=0.497, 95% CI = 0.24-0.75, p<0.001; 932 mitochondrial objects across 10 images, with 5 images per condition), corresponding to approximately 1.64-fold higher ratio than in the control **(Fig. 1C)**. As we used the same microscopy conditions in control and treatment groups, and UV-irradiation was not used to photoconvert mtEOS_green_ in the dataset used by MorphoMapper, an increase in mtEOS_red_ was not expected. Thus, we tested if mtEOS_green_ shows conversion to mtEOS_red_ under different stress conditions that may affect mitochondria, namely osmotic and salt stress conditions **(Fig. S1)**. We found a light-independent increase of mEOS_red_ in presence of 100 mM chloride anions, suggesting that oxidative stress (MtPQ) as well as ionic stress can lead to mEOS_red_ formation in the mitochondrial matrix. As mEOS conversion is linked to a backbone break next to the chromophore, we interpret the increased mitochondrial mEOS_red_ levels as a marker for locally increased protein damage. Thus, matrix mEOS_red_/mEOS_green_ ratio represents a valuable parameter that can be tracked under stress, in addition to its use as a photo-convertible probe. To determine whether the elongated mitochondrial structures after MtPQ treatment reflected increased matrix connectivity, we then irradiated a defined region of interest (ROI), corresponding to approximately 10% of the cell area with an UV diode laser to photoconvert mEOS-labelled mitochondria within that region **(Fig. 1A)**. We found that mitochondrial elongation observed after MtPQ treatment led to faster immediate spread of the mEOS_red_ signal to the ROI comprising the rest of the cell **(Fig. 1D)**. To track mitochondrial elongation over time, this assay was repeated on different sets of cells for five days of treatment with 50 µM MtPQ. MtmEOS_red_ spread increased from the early treatment phase (2-3 h) up to day 3, whereas no significant difference between control and MtPQ-treated cells was present at days 4 and 5 **(Fig. 1D and Fig S2)**. Accordingly, we frequently observed fragmented or doughnut-shaped mitochondria at these later time points **(Fig. S2)**. Visual inspection of the MtPQ-treated protonema culture **(Fig. 1E)** showed partially visible tissue damage and cell bleaching by day 5, whereas control tissue remained largely green and intact. Together, these results show that MtPQ treatment induces transient mitochondrial elongation with higher mitochondrial population connectivity, followed by mitochondrial fragmentation and, in part, leading to cell death.

**Figure 1:**
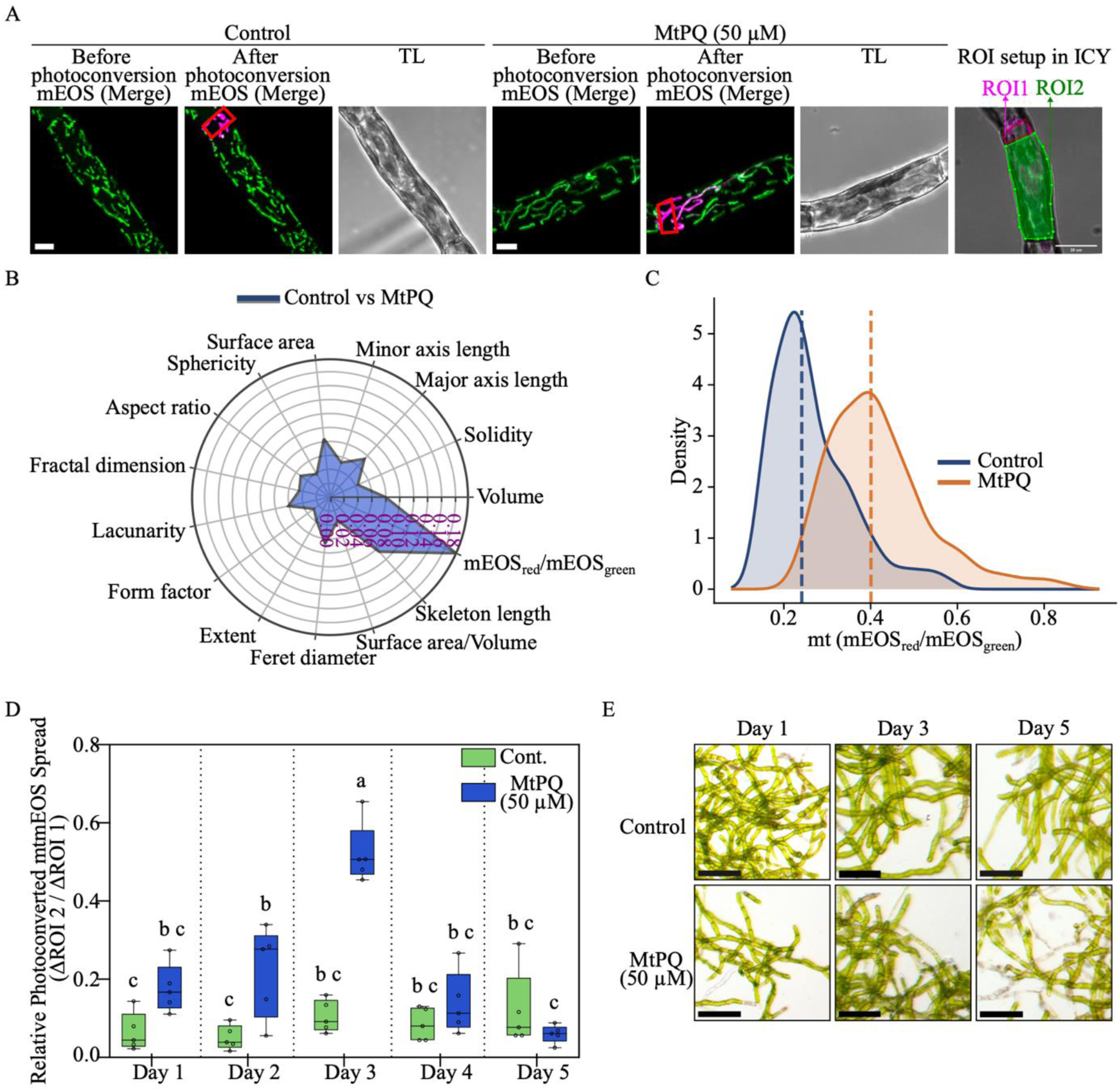
MtPQ induces mitochondrial elongation and alters mitochondrial connectivity in *Physcomitrium patens* protonema. (A) Representative confocal images of *Physcomitrium patens* expressing mitochondrial matrix targeted mtmEOS under control conditions and after treatment with 50 µM MtPQ for 2-3 h. MtPQ-treated cells show visibly elongated mitochondrial structures compared with controls. Green, mtmEOS_green_; magenta, mtmEOS_red_, TL, transmitted light. Scale bar 10 µm. Right panel: representative ROI setup used for quantification of photoconverted mtmEOS_red_ signal spread. ROI1 marks the photoconverted mitochondrial region, and ROI2 marks the non-photoconverted region used to quantify relative spread. Scale bar 20 µm. (B) Feature contribution analysis showing features contributing the most to differences between control and MtPQ-treated samples in the multivariate feature space; feature definitions in Table S2 (C) Density plot of mEOS_red_/mEOS_green_ distribution in control vs. MtPQ-treated cells. Dotted lines represent the median values of 0.24 and 0.40 in control and MtPQ-treated samples, respectively. (D) Quantification of relative photoconverted mtmEOS spread in ROI2 over five days of control and 50 µM MtPQ treatment. Increased spread indicates increased mitochondrial matrix connectivity. Statistical analysis was performed by two-way ANOVA followed by Tukey’s multiple comparison test. Different letters indicate statistically significant differences between groups (*p*<0.0001). (E) Representative stereomicroscopy images of protonema after 1, 3, and 5 days under control conditions or 50 µM MtPQ treatment.

### MtPQ causes shifts in glutathione redox potential in several subcellular compartments

As MtPQ treatment should affect subcellular redox steady states, we tested if the observed changes of mitochondrial morphology and increase in mt mEOS_red_/mEOS_green_ ratio correlated with altered physiological parameters, namely the glutathione redox potential (*E*_GSH_). Here, we used stable transgenic lines expressing the genetically encoded biosensor roGFP2 fused to human glutaredoxin 1 (Grx1) targeted to the cytosol, the chloroplast stroma (Müller-Schüssele *et al*., 2020) as well as the mitochondrial matrix **(Fig. S3)**. Fluorescent moss protonema was analysed after MtPQ treatment in a plate reader-based multi-well setup (Bohle *et al*., 2024; Ugalde *et al*., 2022; Wagner *et al*., 2019) and sensor dynamic range was assessed via an *in vivo* calibration. In the mitochondrial matrix, 5 and 20 µM MtPQ did slightly but not significantly alter the sensor oxidation state (400/482 ratio), whereas 50 µM MtPQ caused an immediate and persistent significant increase **(Fig. 2A, Fig. S4)**. Oxidation of the mitochondrial matrix *E*_GSH_ was substantial, as sensor oxidation nearly reached maximal levels determined via the *in vivo* calibration with hydrogen peroxide. Secondly, we assessed if the MtPQ treatment had effects on the *E*_GSH_ in other subcellular compartments. We found that the cytosolic Grx1-roGFP2 sensor reacted with a transient peak (duration c. 30 min) to MtPQ treatment doses **(Fig. S4)**, followed by a smaller but significant persistent increase in oxidation at 50 µM MtPQ **(Fig. 2B)**. Interestingly, the chloroplast-targeted TK_TP_-Grx1-roGFP2 sensor equally showed a dose-dependent response, with significant persistent oxidation already detected at 20 µM and a stronger oxidation at 50 µM MtPQ **(Fig. 2, Fig. S4)**.

**Figure 2:**
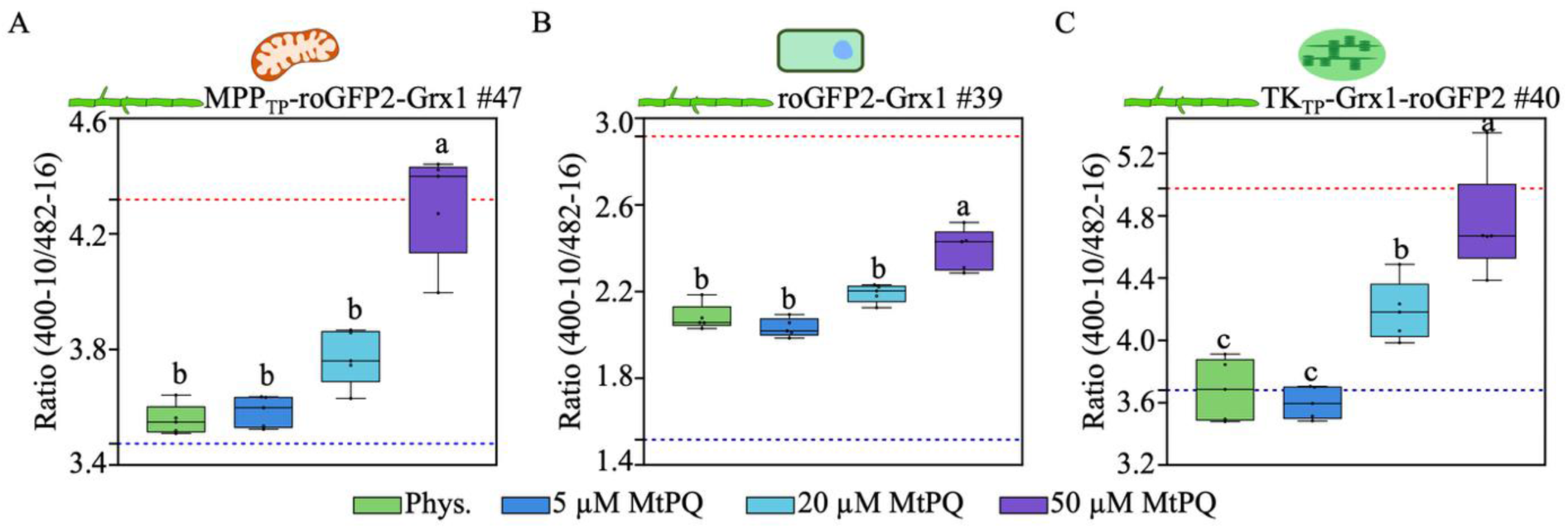
MtPQ dose response of glutathione redox potential *E*_GSH_ in different cellular compartments 2.5 h after adding treatment. (A) Mitochondrial glutathione redox potential using the MPP_TP_-roGFP2-Grx1 #47 sensor after treatment with 5, 20, or 50 µM MtPQ. Only 50 µM MtPQ caused a significant increase in the sensor ratio compared with the physiological control. (B) Cytosolic glutathione redox potential measured using roGFP2-Grx1 #39. A significant increase in the sensor ratio was detected after 50 µM MtPQ treatment. (C) Stromal glutathione redox potential measured using TK_TP_-Grx1-roGFP2 #40. MtPQ induced a dose-dependent increase in the sensor ratio, with significant oxidation observed after 20 and 50 µM MtPQ treatment. Ratios are shown as 400-10/482-16 excitation ratios. Blue and red dashed lines indicate the fully reduced and fully oxidized sensor states after treatment with 10 mM DTT and 10 mM H₂O₂, respectively. Boxes show the interquartile range, horizontal lines indicate medians, whiskers indicate the data range, and individual points represent biological replicates. Different lowercase letters indicate statistically significant differences between treatments, two-way ANOVA followed by Tukey’s post hoc test, *p* < 0.05. Full plate reader time series is shown in Fig. S4.

Thus, MtPQ treatment is not specific for the mitochondrial matrix in *P. patens* but impacts local thiol redox steady states strongest in the mitochondrial matrix, with similar effects on the plastid stroma and, after a transient peak, the lowest effect in the cytosol. In summary, MtPQ-induced oxidative stress causes changes in several local redox steady states that correlate with transient mitochondrial elongation as well as mtmEOS conversion in *P. patens*. This data characterises and connects oxidative stress downstream of superoxide formation by a redox cycler to specific change in mitochondrial morphology, i.e. mainly mitochondrial elongation resulting in higher connectivity. Next, we investigated if changing mitochondrial dynamics in favour of a more connected mitochondrial population would change oxidative stress resilience. To this end, we generated mutants with impaired mitochondrial fission in our established experimental system.

### Loss of PpELM1B affects plant growth, mitochondrial morphology and mtEOS_red/green_ ratio

To create *P. patens* lines with chronically altered mitochondrial dynamics, we targeted the conserved ELM1 (ELONGATED MITOCHONDRIA) protein, that recruits dynamin-related proteins of the mitochondrial fission machinery to plant mitochondria (Arimura *et al*., 2008a; Nagaoka *et al*., 2017). To affect mitochondrial fission, CRISPR/Cas9-based gene editing (Mallett *et al*., 2019) was used to target the two *ELM1* homologs in *P. patens* (*PpELM1A* Pp3c23_3410V3.1 and *PpELM1B* Pp3c20_13230V3.1). Two independent protospacer sequences were designed for each gene, targeting exons two and six **(Fig. 3, Fig. S5)** to create loss-of-function mutations.

**Figure 3:**
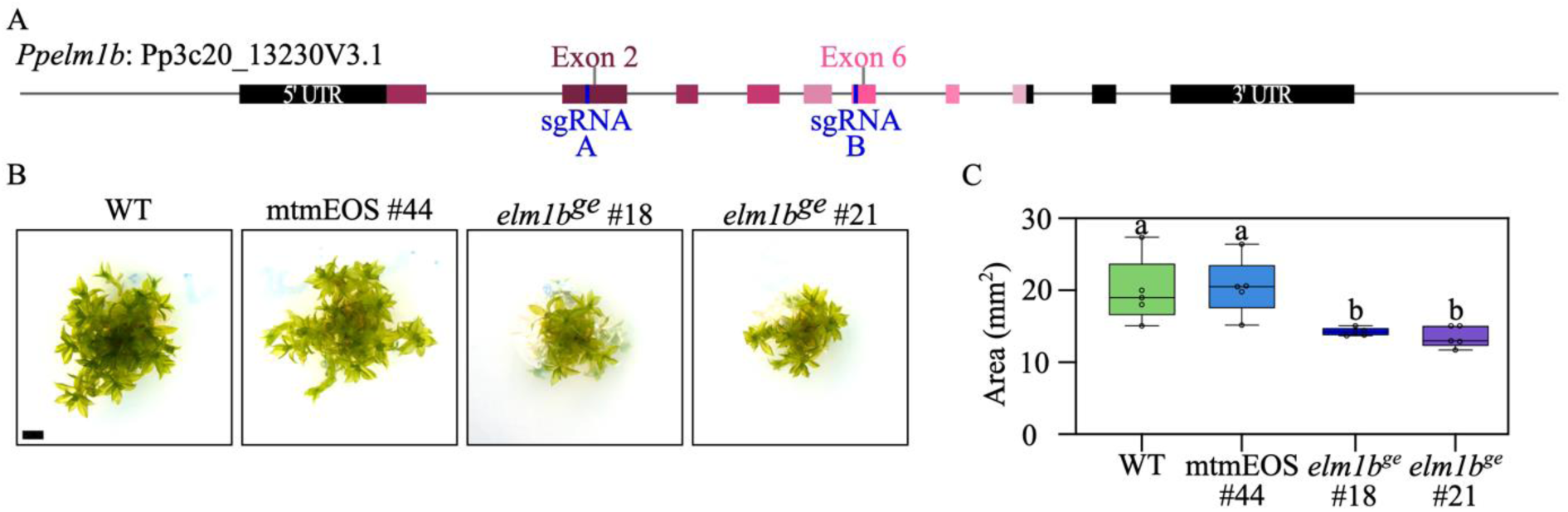
Generation of *elm1b*_ge_ lines in *P. patens*. (A) Schematic representation of the *PpELM1B* gene and CRISPR/Cas9 target sites. (B) Phenotype of two independent *elm1b_ge_ P. patens* lines grown on KNOP ME agar. Scale bar = 1 mm. (C) Quantification of growth. Different letters indicate significant differences between groups. Data represent n = 5 biological replicates. Statistical analysis was performed using one-way ANOVA followed by Tukey’s multiple comparisons test (p < 0.01).

To identify gene-edited lines, PCR-based screening was performed using primer pairs flanking the sgRNA target regions **(Table S1)**. While we did not find stable lines with mutations in *PpELM1A* among candidate plants surviving selection, we were able to obtain *Ppelm1b* gene-edited lines in both fluorescent reporter background lines mtmEOS#44 and MPP_TP_-roGFP2-Grx1#47. PCR screening detected shorter fragments than the wildtype (WT) allele (**Figs. S5, S6**) and sequencing confirmed mutations causing frameshifts and premature stop codons in exon 2 **(Figs. S5, S6)**. Mutant line gametophore development was normal, but overall growth was slower than WT and the mtmEOS#44 background line **(Fig. 3B, C)**.

To investigate whether the loss of *Pp*ELM1B affects mitochondrial morphology, confocal fluorescence imaging was performed in two independent *Ppelm1b^ge^* lines (#18 and #21) and in their mtmEOS #44 genetic background. In both gene-edited lines mitochondria appeared visibly more elongated and interconnected. Additionally, we observed a signal increase in the mtmEOS_red_ channel **(Fig. 4A)**. We compared mitochondrial morphology between mtmEOS#44, MtPQ-stressed mtmEOS#44 and *Ppelm1b^ge^* lines using the MorphoMapper tool. Non-Metric Multidimensional Scaling (NMDS) analysis yielded four clusters, with the clusters of the *Ppelm1b^ge^* lines largely overlapping, indicating similarity **(Fig. 4B)**. The main contributions to the dissimilarity between mtmEOS#44 and *Ppelm1b^ge^*lines were the mEOS_red_/mEOS_green_ ratio, follow by ‘lacunarity’ **(Fig. 4C)**. Notably, chronic mitochondrial morphology changes in *Ppelm1b^ge^*lines were distinct from transient mitochondrial elongation in MtPQ-stressed mtmEOS#44, with ‘volume’, ‘skeleton length’, ‘surface area’ contributing most to the dissimilarity of morphologies **(Fig. 4D**, feature definitions in **Table S2)**. Total mitochondria volume per cell was significantly increased whereas the total number of mitochondria decreased in *Ppelm1b^ge^* lines compared to the mtmEOS#44 background **(Fig. S7)**. Thus, mitochondria elongate in both MtPQ treatment as well as in absence of ELM1B but show also distinguishable and specific morphological changes.

**Figure 4:**
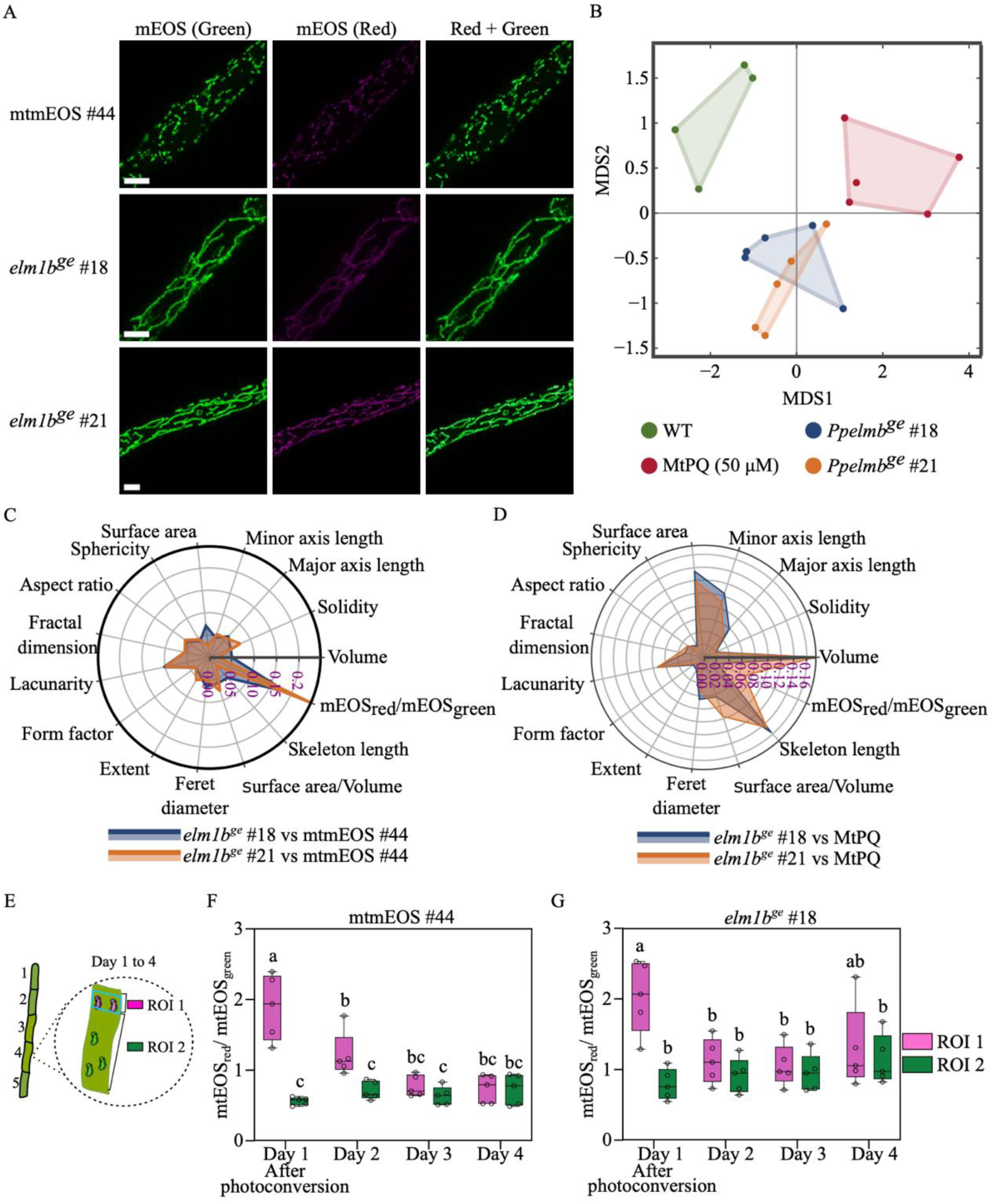
Mitochondrial morphology of *elm1b*_ge_ lines in *P. patens*. (A) Maximum intensity projections (MIPs) of confocal z-stacks showing mitochondrial morphology in the WT control (mtmEOS #44) and two *Ppelm1B_ge_* lines (#18 and #21). Channels represent before mEOS conversion in green, photoconverted red mEOS in magenta, and merged signals. Scale bars = 10 µm. (B) Multidimensional scaling (MDS) embedding of mitochondrial population morphology across different groups. Each point represents one image. The *Ppelm1b_ge_* samples were positioned close to each other, while WT and MitoPQ samples occupied distinct regions of the MDS space. (C, D) Feature contribution analysis to identify the features that contribute the most to separation of two *Ppelm1b_ge_* vs the control (mtmEOS #44) (C) and the two *Ppelm1b_ge_* lines from MtPQ treatments (D). (E) Schematic overview of cell and ROI selection in embedded protonemal tissue. On day 1, approximately 10% of the cell area was photoconverted and the same cell was imaged daily for four days. ROI 1 indicates the initially photoconverted region, whereas ROI 2 indicates the non-photoconverted region. (F, G) Quantification of mEOS red-to-green fluorescence ratios in ROI 1 and ROI 2 in mtmEOS #44 (F) and *elm1b_ge_* #18 (G). Data are shown for n=5 cells. Statistical significance was tested by two-way ANOVA followed by Tukey’s multiple comparisons test. Different letters indicate statistically significant differences between groups, p < 0.0001.

As mixing of mitochondrial matrix content may be affected by impairing fission, we followed cells with partially photoconverted mitochondria over a longer time course. As the observed mixing of mitochondrial matrix content in *P. patens* was slow, we embedded protonema filaments in low-melting point agar and repeated confocal microscopy of the same photoconverted cells daily **(Fig. 4E)**. The mitochondrial mEOS_red_/mEOS_green_ ratio was successfully significantly shifted by photoconversion in ROI1 and subsequently decreased, until it was statistically indistinguishable from the initially non-photoconverted rest of the cell (ROI2) on the third day after photoconversion **(Fig.4F; Figs. S8 and S9)**. These data show that mixing of matrix content takes place in *P. patens* and that the time span for homogenisation the mitochondria population of a cell lies in the range of days. Notably, mt mEOS_red_/mEOS_green_ ratios in the *Ppelm1b^ge^* cells were already converging on the second day after partial photobleaching, indicating persistent matrix mixing and faster homogenisation of mitochondrial matrix content **(Fig. 4F,G; Figs. S8 and S9)**.

### Loss of PpELM1B leads to a less negative matrix *E*_GSH_ and reduced dark respiration

To investigate if the observed and characterised chronic changes to mitochondrial population dynamics lead to physiological changes, we used ELM1B gene-edited mutants in the moss line expressing matrix-targeted roGFP2-Grx1 **(Fig. S6)**. To compare sensor oxidation between mutants and background plant lines, we determined the degree of oxidation (OxD) of mitochondrial roGFP2-Grx1 by performing *in vivo* sensor calibration, again using a plate reader-based set up and protonema cultures **(Fig. 5**, full time series **Fig. S10)**. By comparing sensor oxidation under physiological conditions to the values obtained by calibration with either 10 mM DTT or 10 mM H_2_O_2_, we found an OxD_roGFP2_ of 25 +/-7% in the MPP_TP_-roGFP2-Grx1#47 background line compared to 97 +/-0.5% and 99 +/-5% in *Ppelm1b^ge^* #8 and *Ppelm1b^ge^* #26, respectively **(Fig. 5A)**. This indicates that mitochondrial roGFP2-Grx1 in the *Ppelm1b^ge^* lines is nearly completely oxidized without additional oxidative treatment under the tested physiological conditions. Based on a matrix pH of 7.8 (Schwarzländer *et al*., 2008), the resulting calculated matrix *E*_GSH_ is c. -318 mV in the MPP_TP_-roGFP2-Grx1 #47 background line, compared to an oxidative shift of approximately 60-80 mV in *Ppelm1^ge^* lines with an *E*_GSH_ of c. -259 mV for *Ppelm1b^ge^* #8 and *E*_GSH_ c. -237 mV for *Ppelm1b^ge^* #26. As some cells did show weak cytosolic/nuclear roGFP2-Grx1 signal or increased autofluorescence, parameters that cannot be addressed by a plate reader-based setup, we additionally assessed the redox state of matrix-localised roGFP2-Grx1 by confocal microscopy using mitochondrial regions of interest for ratiometric analysis **(Fig. S11)**. Microscopic analysis confirmed that matrix-localised roGFP2-Grx1 is nearly fully reduced in the sensor line, while both *Ppelm1b^ge^*lines in that genetic background show nearly full sensor oxidation **(Fig. S11)**. Thus, we find that impaired mitochondrial fission in *P. patens* causes a substantially less reducing matrix glutathione redox potential.

**Figure 5:**
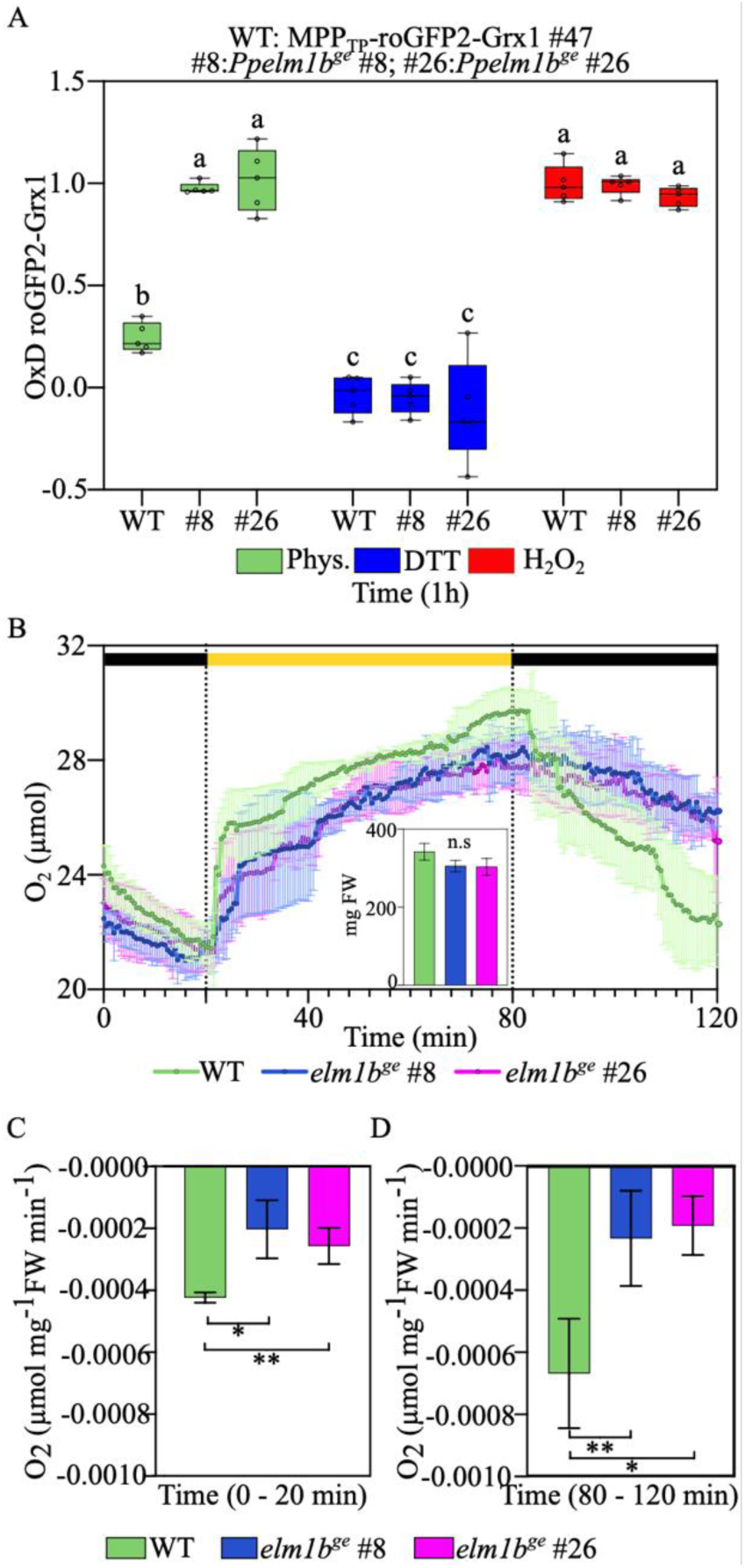
Loss of PpELM1B alters the mitochondrial glutathione redox state and respiration in *P. patens*. (A) Degree of oxidation (OxD) of mitochondrial roGFP2-Grx1 in the WT reporter line MPP_TP_-roGFP2-Grx1 #47 and the two independent *Ppelm1b_ge_* lines (#8, #26) under physiological conditions or following treatment with 10 mM DTT or 10 mM H_2_O_2_. Data correspond to plateau values after 1h of plate reader-based read-out (Fig. S10). Box plots show the median, the 25th–75th percentiles, and the minimum and maximum values (n = 5). Different letters indicate statistically significant differences between groups, determined by two-way ANOVA followed by Tukey’s multiple-comparisons test (P < 0.0001). (B) Available dissolved O_2_ concentration (µmol) in liquid cultures of WT, *Ppelm1b_ge_* #8 and *Ppelm1b_ge_* #26 during a dark-light-dark cycle. Cultures were pre-incubated in dark for 2 h and then measured for 20 min in dark, followed by 60 min of illumination at 100 µmol photons m_-2_s_-1_ and another 40 min dark period. Black and yellow bars indicate dark and light periods, respectively. The inset shows the fresh weight (FW) of the samples after the oxygen measurements. Lines show the mean and error bars represent SD (n = 3). (C, D) Rates of O_2_ change, normalised to fresh weight (FW), during the initial dark period from 0-20 min (C) and the final dark period from 80-120 min (D). Data are shown as mean ± SD. Statistical significance was determined using pairwise unpaired two-tailed *t*-tests; p < 0.05 (*) and p < 0.01 (**).

As the altered mitochondrial morphology and matrix glutathione redox steady state in the *Ppelm1b^ge^* lines suggested that mitochondrial function may also be affected, we next assessed respiration. We measured oxygen concentration in parallel protonema cultures with adjusted culture density of WT and two independent *Ppelm1b^ge^* lines (#8 and #26) in a dark/light/dark regime using a Clark-type oxygen microsensor with a 200 µm tip diameter inserted into a permanently stirred protonema culture. Before starting the measurement, 100 ml protonema cultures were incubated in the dark for 2 h to allow respiration rates to reach a steady state condition. The measurement then included an initial dark phase for 20 min, a light phase for 60 min, and a final dark phase for 40 min. In the dark, oxygen levels decreased steadily, reflecting oxygen consumption, whereas levels rapidly increased in the light in response to active photosynthesis. WT cultures showed a reproducible stronger decline in oxygen concentration in the dark than the *Ppelm1b^ge^* lines **(Fig. 5B)**. Thus, respiratory activity in *Ppelm1b^ge^* lines is decreased.

### *elm1b^ge^* lines are more sensitive to oxidative stress

As impaired fission impacted mitochondrial morphology and physiology, we finally tested the effect of additional oxidative stress on *Ppelm1b^ge^*lines on a microscopic as well as macroscopic level. Investigating MtPQ-treated *Ppelm1b^ge^* lines microscopically, we found that the protonema cells treated with 50 µM MtPQ showed more pronounced changes in mitochondrial morphology in *Ppelm1b^ge^* #18 line than in the respective mtmEOS #44 background line **(Fig. S12)**. Treatment of density-adjusted parallel protonema cultures with 50 µM MtPQ resulted in visible bleaching of the MtPQ-treated *Ppelm1b^ge^* cultures by day 5 **(Fig. 6)**, indicating increased sensitivity to additional oxidative stress.

**Figure 6:**
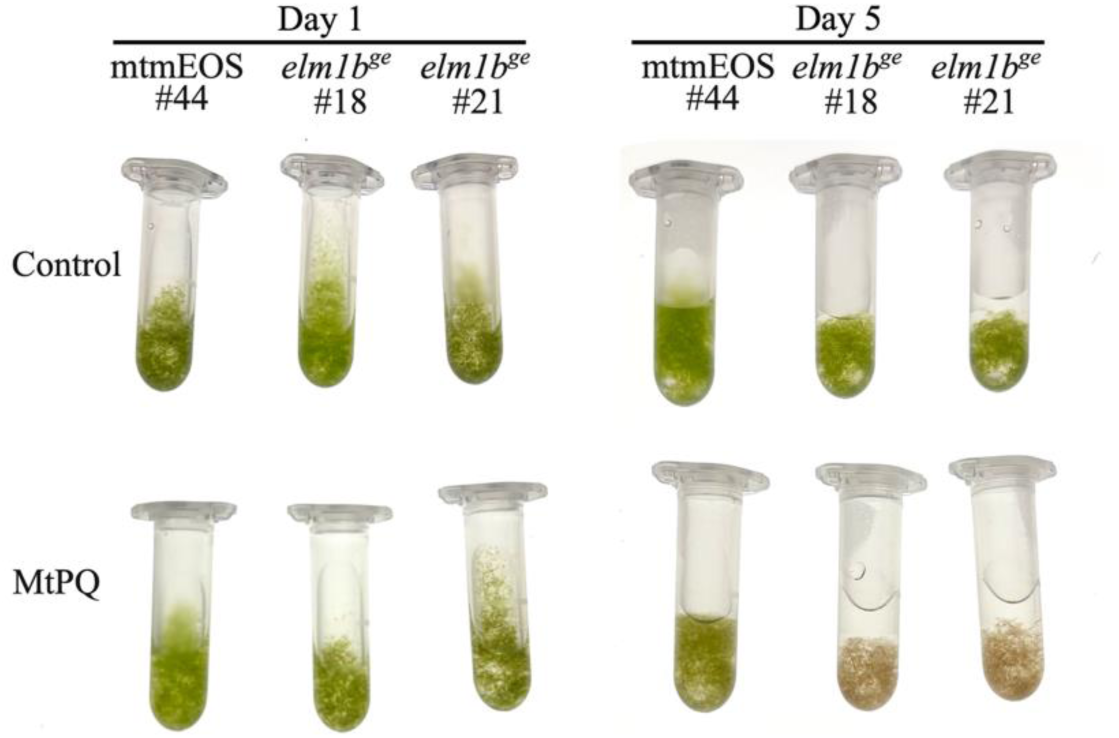
*Ppelm1b*_ge_ lines are more sensitive to oxidative stress. Loss of PpELM1B increases susceptibility to mitochondrial oxidative stress induced by MtPQ. Representative images of liquid cultures of the mtmEOS #44 background line and two independent *Ppelm1b_ge_* lines, under control conditions and after MtPQ treatment. Images were taken on day 1 after 2 h of incubation and on day 5 after prolonged incubation with the stressor.

To track if the observed effect was compartment or oxidant-specific, we additionally tested treatments with the thiol-specific oxidant DPS (2,2’-dipyridyl disulfide) and exogenous addition of 10mM H_2_O_2_ **(Figs. S13, S14)**. We found that *Ppelm1b^ge^* lines macroscopically showed not difference to the MPP_TP_-roGFP2-Grx1 background after 10mM H_2_O_2_ treatment while they exhibited increased sensitivity to 0.5mM and higher concentrations of DPS. These results indicate that loss of PpELM1B is associated with increased sensitivity to oxidative stress, as induced by MtPQ, as well as to direct thiol oxidation.

## DISCUSSION

### Oxidative stress affects organelle morphology, heterogeneity and physiology

Oxidative stress affects organelle physiology and triggers a cellular stress response that may re-configure mitochondrial network morphology. We tested how the mitochondrial population responds to oxidative stress in a non-seed plant model with high amenability to microscopic analyses, *P. patens*. Paraquat (methyl viologen) is a herbicide with known oxidative effects for both mitochondria and chloroplasts in plants, with the main effect, that is light-dependent, in chloroplasts (Ugalde *et al*., 2021; Vicente *et al*., 2001; Halliwell and Gutteridge, 2015): Paraquat is amplifying superoxide generation as a redox cycling agent accepting electrons from Fe-S proteins at photosystem I as well as flavoproteins, including complex I, generating superoxide at the matrix side (K., Khan *et al*., 2024; Halliwell and Gutteridge, 2015). MtPQ was designed as paraquat conjugated to the mitochondria-targeting triphenylphosphonium cation for membrane potential-dependent specific import and local boosting of superoxide production in mitochondria (Robb *et al*., 2015). As tested with genetically encoded redox sensors in *Arabidopsis thaliana*, MtPQ-induced stress can be dosed to affect the matrix but not the cytosol (K., Khan *et al*., 2024). Using glutaredoxin-linked roGFP2 biosensors that respond to changes in the glutathione redox potential *E*_GSH_ (Gutscher *et al*., 2008; Schwarzländer *et al*., 2016; Müller-Schüssele *et al*., 2021; Meyer and Dick, 2010), we found an immediate oxidative response to MtPQ in the chloroplast stroma, the cytosol and the mitochondrial matrix, using a plate reader-based readout. In our hands, 50 µM MtPQ was required to induce a significant oxidative shift for mitochondrial roGFP2-Grx1 excitation ratio in *P. patens*. However, the same treatment also affected sensor oxidation states of cytosol and stroma, indicating that the MtPQ-induced stress response was not confined to the mitochondrial matrix in *P. patens*. Thus, in *P. patens* as experimental system, it is possible to induce matrix oxidation with MtPQ, however with parallel stroma oxidation and a lower but significant oxidative shift in the cytosol. It is currently not possible to conclude what process triggers stroma and cytosolic oxidation as paraquat has several possible electron donors. Moreover, MtPQ treatment was conducted in the relative dark inside a plate reader, where light-dependent superoxide formation at PSI by PQ was limited using *A. thaliana* (Ugalde *et al*., 2021). However, H_2_O_2_ generated downstream of paraquat-dependent superoxide formation at PSI can affect cytosolic *E*_GSH_ (Ugalde *et al*., 2021) and we cannot exclude that also mitochondrial H_2_O_2_ might affect redox processes in other subcellular compartments (Noctor and Foyer, 2016; Huang *et al*., 2016; Schwarzländer and Finkemeier, 2013). This redox link between compartments likely involves local superoxide detoxification, H_2_O_2_ diffusion and local activities of enzymes scavenging peroxides, such as the ascorbate glutathione cycle enzyme dehydroascorbate reductase that draws electrons from the glutathione pool, generating glutathione disulfide (Ugalde *et al*., 2021; Rahantaniaina *et al*., 2017; Waszczak *et al*., 2018). Thus, on the one hand, MtPQ is a useful tool to rapidly manipulate redox steady state of the mitochondrial matrix without blocking the respiratory chain. On the other hand, its use in plants, that also possess a second compartment that might be influenced by PQ, requires careful evaluation of results. Here, genetically encoded redox sensors are vital to track local and dynamic effects of stress treatments. In summary, the MtPQ treatment in *P. patens* is not suitable to distinguish between the potential origins of oxidative stress nor downstream signalling events but generates an oxidative stress that affects mainly the endosymbiont-derived organelles **(Fig. 7** left panel).

**Figure 7:**
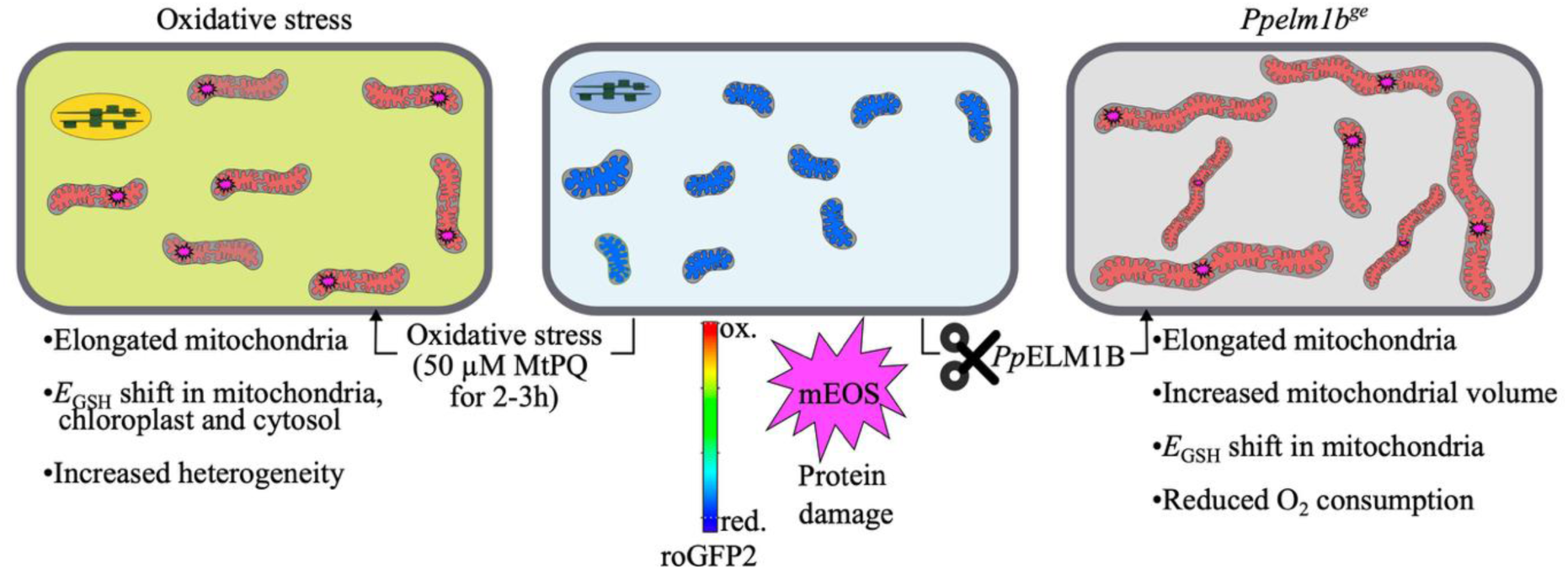
Linking populations dynamics and physiological parameters. Treatment with 50 µM MtPQ induced mitochondrial elongation, oxidation of mitochondrial, cytosolic and chloroplast-targeted Grx-linked roGFP2, and a broader distribution of mitochondrial mEOS red-to-green ratios. Loss of PpELM1B was associated with fewer and more elongated mitochondria, increased total mitochondrial volume per cell **(Fig. S7)**, increased oxidation of the mitochondrial roGFP2-Grx1 sensor, higher mitochondrial mEOS red-to-green ratios and reduced respiration in darkness. Loss of PpELM1B also increased sensitivity to prolonged MtPQ treatment and DPS-induced thiol oxidation.

Using 50 µM MtPQ in *P. patens*, oxidative stress was sufficiently severe to cause macroscopically observable effects but allowed cell survival, which enabled us to study the mitochondrial population response during several days. Here, we show that the mitochondrial population in *P. patens* responds to this oxidative stress by elongation that increases matrix connectivity within a cell by a higher average individual volume of fewer elongated mitochondria. While the mitochondrial population was more connected during several days of withstanding oxidative stress, matrix mixing in *P. patens* was not fast enough to lead to complete homogenisation of the population, as evidenced by increased heterogeneity of observed mEOS_red_/mEOS_green_ ratio in the individually segmented mitochondria **(Fig. 7** left panel). Mitochondrial hyperfusion is already known as a mitochondrial stress response in animal and fungal systems (Friedman and Nunnari, 2014), where a link to protection from autophagy and maintenance of ATP production capacity was evidenced (Liesa and Shirihai, 2013). To date, in plant systems, mitochondrial hyperfusion was characterised prior to cell division with possible links to homogenisation of mtDNA content and distribution (Sheahan *et al*., 2005; Seguí-Simarro *et al*., 2008; Arimura *et al*., 2004). Fusion and transient ‘kiss-and-run’ encounters (Liu *et al*., 2009) permit matrix-content exchange in the multipartite plant mitochondria population: previous investigations using photoconvertible fluorescent proteins have shown high fusion and fission rates, with complete matrix mixing in one cell occurring as fast as within an hour (Arimura, 2018; Arimura *et al*., 2004). Therefore, in plants, fusion fission dynamics supporting a ‘social’ mitochondrial network with frequent encounters and exchange has been evidenced (Chustecki *et al*., 2021; Giannakis *et al*., 2022). In *P. patens*, prolonged exposure to oxidative stress led to mitochondrial fragmentation including doughnut-shapes, features that have been associated with (reversible) loss of functionality and damage in animal systems (Ahmad *et al*., 2013; Liu and Hajnóczky, 2011).

In summary, we interpret the dynamic changes to mitochondrial morphology under oxidative stress in *P. patens* as a population response supporting survival, potentially by sharing stress, minimizing stress-induced heterogeneity and maintaining mitochondrial functions.

### Mitochondrial fission safeguards oxidative stress resilience

As fusion/fission dynamics underpin changes in the mitochondrial matrix exchange and morphology, we investigated the effect of chronically elongated mitochondria on mitochondrial performance and overall plant stress resilience. Mitochondrial fusion is currently not mechanistically understood in plants: mitofusin/FZL function is not conserved (Gao *et al*., 2006), while several non-homologous factors influencing mitochondrial fusion have been evidenced (Kenneally *et al*., 2026; White *et al*., 2020; El Zawily *et al*., 2014). Thus, we chose to create null mutants of the mitochondria-specific fission factor ELM1 that recruits evolutionary conserved dynamin-related proteins (DRP3 gene family in plants) to division sites (Arimura *et al*., 2008b; Fujimoto *et al*., 2009; Nagaoka *et al*., 2017). According to models, plant mitochondrial fission is vital for the exchange of mitochondrial content across the organelle population (Chustecki *et al*., 2021; Johnston, 2019). Additionally, first experimental evidence links fission to segregation of damaged mitochondrial content via mitophagy (Broda *et al*., 2018; Mao *et al*., 2013; Pettinari *et al*., 2022; Ren *et al*., 2021; Ma et al., 2025), similar to animal and fungal cells (Wikstrom *et al*., 2009; Twig *et al*., 2008; Labbé *et al*., 2014; Abeliovich *et al*., 2013). As in *A. thaliana* and in the liverwort *Marchantia polymorpha*, null mutants of *P. patens* ELM1B were viable and showed elongated mitochondria (Arimura *et al*., 2008b; Nagaoka *et al*., 2017). In our hands, we were not able to retrieve single mutants of the second homolog *Pp*ELM1A or double mutants, potentially indicating a vital role for *Pp*ELM1A. However, *Pp*ELM1B is necessary for maintaining normal mitochondrial fission and mitochondrial function, as we found reduced plant growth and respiration rates. The elongated mitochondria in *Ppelm1b^ge^* lines exhibited potentially increased protein damage levels, as evidenced microscopically by increase of mEOS_red_, and less reducing matrix *E*_GSH_, as evidenced by increased roGFP2-Grx1 oxidation levels **(Fig. 7** right panel). As the observed effects in null mutants are chronic, we cannot conclude whether increased thiol oxidation is cause or effect of the observed other phenotypes. We speculate that decreased mitochondrial fission might lead to accumulation of non-functional mitochondrial proteins, eventually leading to increased ROS formation and permanently increased thiol oxidation, that cannot be compensated. A chronically shifted matrix *E*_GSH_ might in turn influence redox steady states of mitochondrial thiol switches (Bohle *et al*., 2024; Nietzel *et al*., 2020; Møller *et al*., 2020), perturbing metabolic regulation. The observed sensitivity of *Ppelm1b^ge^* lines to the specific thiol oxidant DPS (Lopez-Mirabal *et al*., 2007) as well as to MtPQ-induced oxidative stress supports the hypothesis that thiol redox plays a central role in the observed defects. Taken together, our current model links mitochondrial fission in plants to oxidative stress resilience and suggests a link between fission, protein damage levels and thiol redox steady state.

Notably, our analysis revealed differences between MtPQ-induced oxidative stress and ELM1B deficiency that became only apparent when considering the full mitochondrial population by automated 3D-segmentation and feature mapping (MorphoMapper). Although both investigated conditions were associated with mitochondrial elongation and a reduced mitochondrial number, their population-level morphological distributions remained clearly distinct, e.g. in the extent of increased mtmEOS_red_.

Using full photoconversion of mtmEOS_green_ and matrix homogenization analyses over several days, we found that the overall matrix mixing rate in the mitochondrial population of a *Ppelm1b^ge^* cell was not reduced, but rather increased. This indicates that mitochondrial fusion in combination with a higher volume of individual mitochondria was sufficient to maintain matrix protein exchange within the mitochondrial network. However, we found that homogenization of matrix content within one cell takes days in *P. patens*, as opposed to hours in flowering plants. This correlates with the lower movement velocities of mitochondria in *P. patens* (Furt *et al*., 2012). Our data raises the question if the fast matrix mixing observed in flowering plants is an evolutionarily derived feature. Notably, mtDNA complexity and RNA editing sites increased during land plant evolution (Small *et al*., 2020; Gualberto and Newton, 2017; Knoop, 2004) with both factors constituting potential reasons requiring fast and constitutive exchange of mitochondrial content. This is congruent with the fact that some proteins such as editing factors have been found at a sub-stochiometric amount (i.e. less than one molecule in average per mitochondrion) in *A. thaliana* (Fuchs *et al*., 2020). The future challenge includes to visualise and characterise mtDNA and mitochondrial protein exchange in additional model plants and to further understand the mechanistic reasons for the structure and behaviour of plant mitochondrial populations.

### Supplementary data

The following supplementary figures and tables are available:

***Fig. S1***: Confocal microscopy of mitochondria-targeted mEOS under different abiotic stress conditions in *P. patens*.

***Fig. S2***: MtPQ treatment increases mitochondrial connectivity.

***Fig. S3***: Confocal microscopy confirms mitochondrial localization of mitochondria-targeted roGFP2 in *P. patens*.

***Fig. S4***: Dose-dependent oxidation of roGFP2-Grx1 sensors by MtPQ in different subcellular compartments.

***Fig. S5***: Generation and screening of CRISPR/Cas9-mediated *Ppelm1b^ge^* lines in the mtmEOS #44 background.

***Fig. S6***: Generation and screening of CRISPR/Cas9-mediated *Ppelm1b^ge^* lines in the MPP_TP_-roGFP2-Grx1 #47 background.

***Fig. S7***: Loss of ELM1B alters total mitochondrial volume and number in *P. patens*.

***Fig. S8***: Replicate images of mEOS signal redistribution on day 2 after photoconversion.

***Fig. S9***: Replicate images of mtEOS signal redistribution on day 3 after photoconversion.

***Fig. S10***: Plate reader-based comparison of *Ppelm1b^ge^*lines with the mitochondria-targeted roGFP2-Grx1-expressing background.

***Fig. S11***: Calibration of mitochondria-targeted roGFP2-Grx1 in *P. patens*.

***Fig. S12***: Representative confocal images of mtmEOS #44 background line and the

*Ppelm1b^ge^*#18 line with 50 µM MtPQ.

***Fig. S13***: *Ppelm1b^ge^* lines show increased sensitivity to DPS treatment.

***Fig. S14***: *Ppelm1b^ge^* lines did not show visible sensitivity to H_2_O_2_ treatment.

***Table S1***: Oligonucleotides.

***Table S2.*** Shape descriptors used for mitochondrial profiling.

## Acknowledgements

We thank Aurora Martin, Alexander Nies, Marie Jagemann (Molecular Botany, RPTU) and PD Dr. Michelle Gehringer (Microbiology, RPTU) for assistance. We are grateful to Dr. Oliver Trentmann for useful discussions.

## Author Contributions

SST and SJMS designed the research. SP, CG, IN, SST performed experiments and analysed data. ST analysed and curated data. TM and SJMS supervised the research and provided resources. SST and SJMS wrote the manuscript with contributions from all authors. All authors approved the manuscript before submission.

