## Supplemental Information for "Loss of ELM1B impairs mitochondrial fission, matrix redox state and stress tolerance in *Physcomitrium patens*"

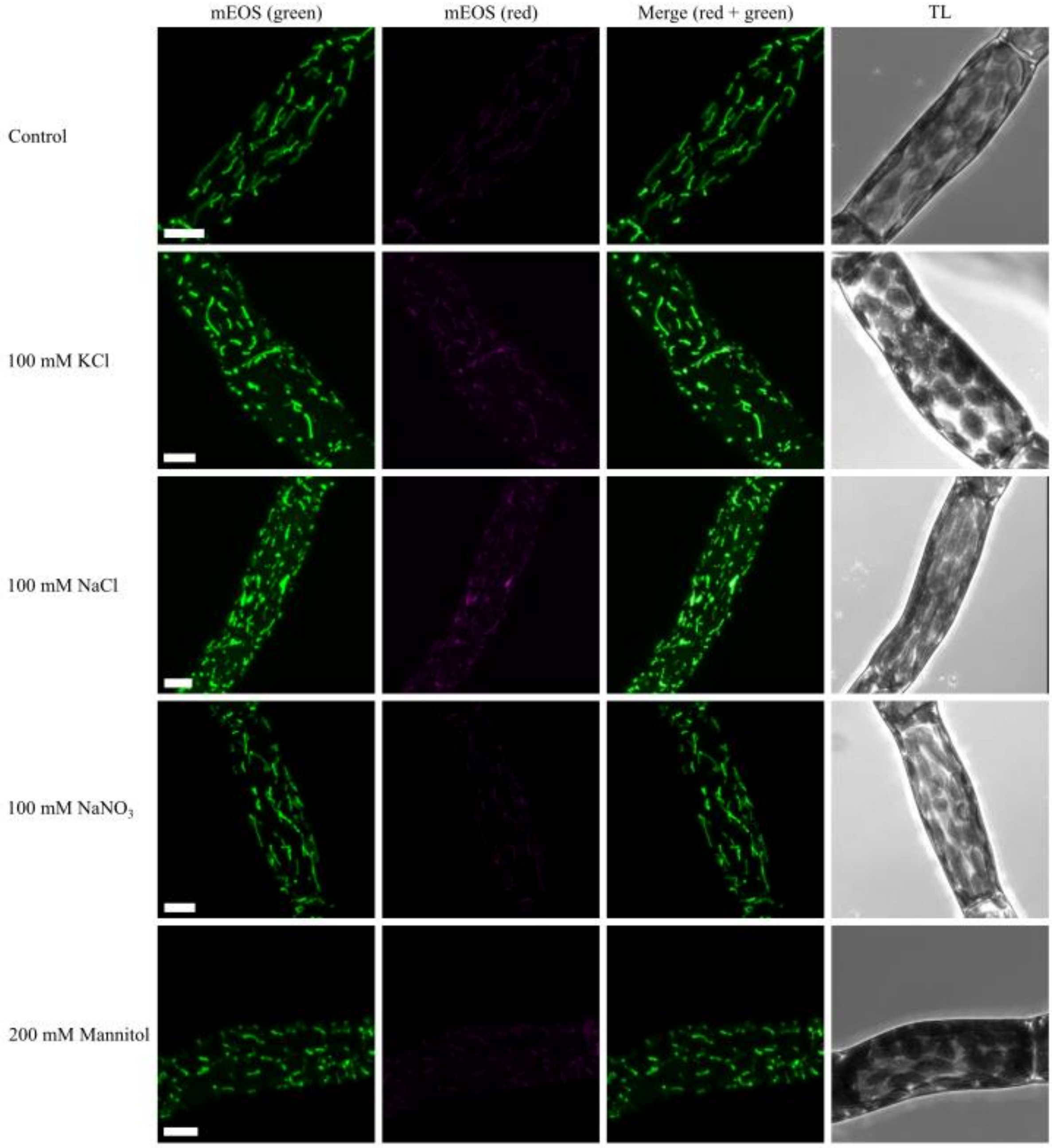


**Supplementary Figure S1: Confocal microscopy of mitochondria-targeted mEOS under different abiotic stress conditions in *P. patens.***
Ionic stress induces mEOS conversion in the mitochondria-targeted mtmEOS in *P. patens*. Representative confocal images of protonema cells treated with 100 mM KCl, 100 mM NaCl, 100 mM $NaNO_3$, or 200 mM mannitol for 2 h in the dark. Channels represent $mEOS_{green}$ (green), $mEOS_{red}$ (magenta) and merged signals as well as transmitted laser light (TL). In the absence of UV irradiation, KCl- and NaCl-treated cells showed a stronger $mEOS_{red}$ signal compared to control cells. Scale bars: 10 µm.

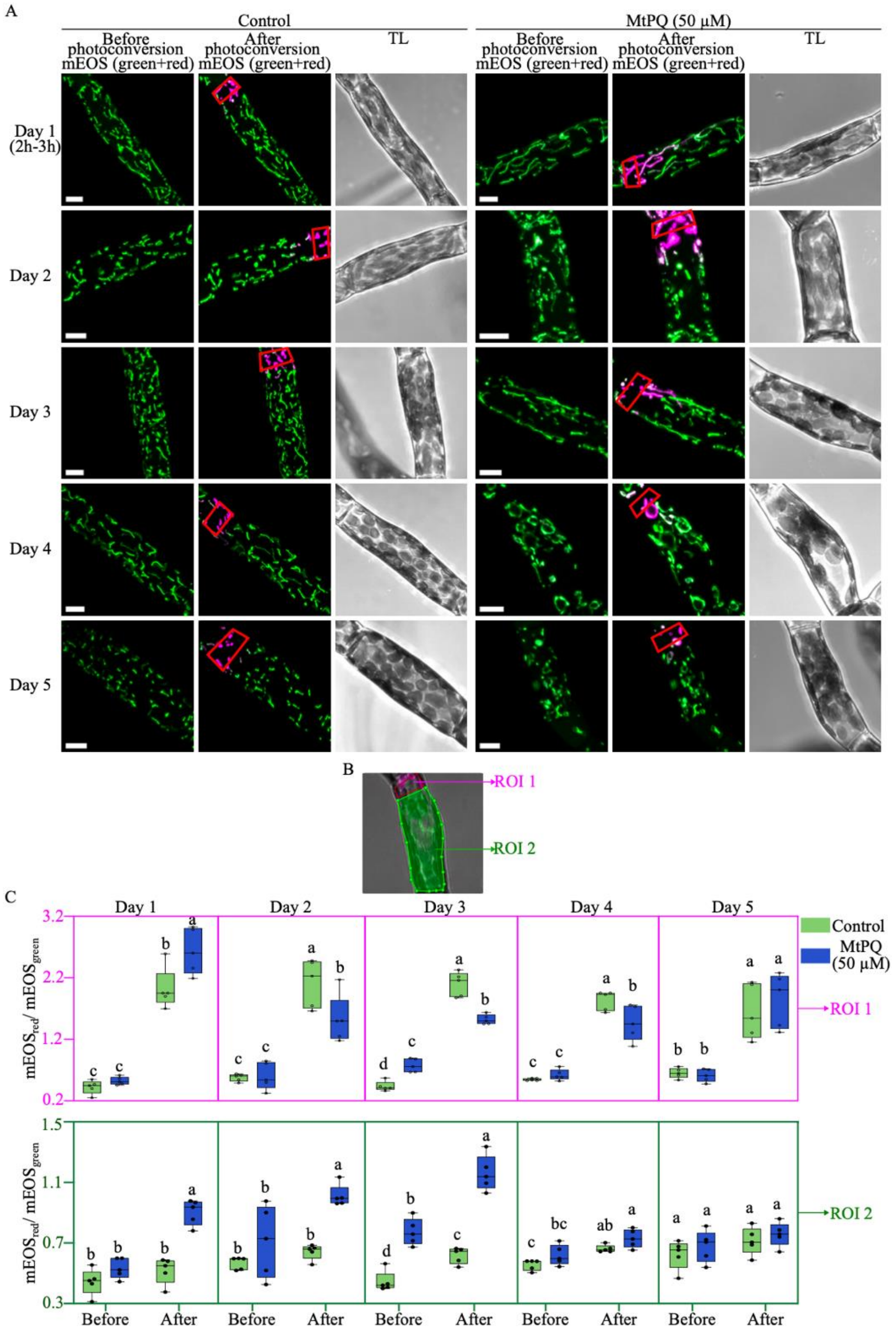


**Supplementary Figure S2: MtPQ treatment increases mitochondrial connectivity**.

MtPQ treatment increases the spreading of photoconverted mitochondrial mEOS signal in *P. patens* protonema cells immediately after photoconversion during the three initial days of treatment. **A** Representative confocal image of protonema cells expressing mitochondria-targeted mtmEOS under control conditions and after treatment with 50 µM MtPQ. For each condition, 5 mL cultures were maintained under standard growth conditions, and 1 mL culture was taken for imaging on each day. A defined region corresponding to approximately 10% of the cell was photoconverted. Maximum intensity projections (MIPs) from confocal z-stacks are shown. Merged images show $mEOS_{green}$ in green and photoconverted $mEOS_{red}$ in magenta. Transmitted light (TL) images are shown for reference. Scale bars = 10 µm. **B** Quantification of the $mEOS_{red}/mEOS_{green}$ fluorescence intensity ratio in two regions of interest before and after photoconversion from day 1 to day 5 of 50 µM MtPQ treatment. ROI 1 corresponds to the photoconverted region, and ROI 2 corresponds to the non-photoconverted region of the same cell. Data are shown for n = 5 cells per day and condition. Statistical analysis was performed separately for each day and ROI using ordinary two-way ANOVA with treatment (control vs. 50 µM MtPQ) and photoconversion status (before vs. after photoconversion) as factors, followed by Tukey's multiple comparisons test, $p < 0.05$.

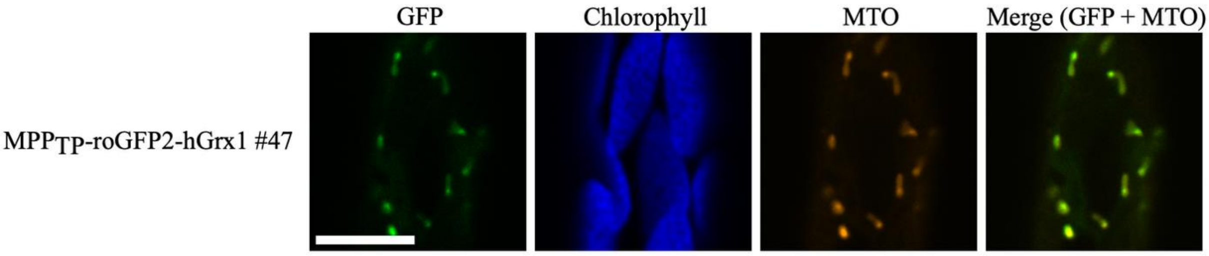


**Supplementary Figure S3: Confocal microscopy confirms mitochondrial localization of mitochondria-targeted roGFP2 in *P. patens*.**
Representative confocal microscopy images of protonema cells expressing mitochondria-targeted roGFP2-Grx1 (MPP$_{TP}$-roGFP2-Grx1 #47). Protonema cultures were incubated with MitoTracker Orange (MTO) in KNOP ME medium (1 nM final concentration) for 15 min before imaging. The roGFP2 fluorescence signal was excited with a 488 nm laser and is shown in green. MTO was excited with a 543 nm laser and is shown in orange. Chlorophyll autofluorescence is shown in blue. Emission was detected at 507–536 nm for roGFP2 fluorescence, 580–624 nm for MTO and 657–687 nm for chlorophyll autofluorescence. Merged images show the overlap between roGFP2 and MTO signals. Scale bar = 10 µm.

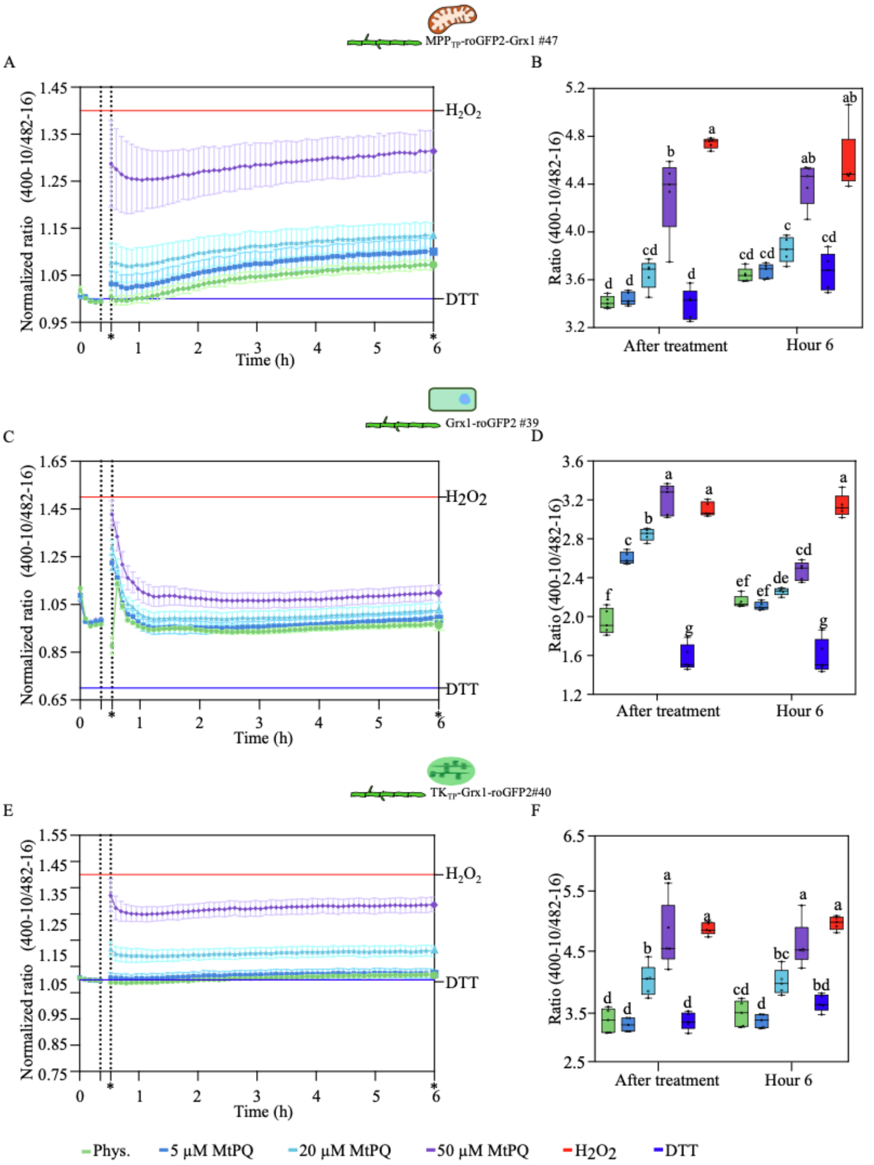


**Supplementary Figure S4: Dose-dependent oxidation of roGFP2-Grx1 sensors by MtPQ in different subcellular compartments.**

MtPQ induces a dose-dependent increase in roGFP2-Grx1 sensor oxidation in the mitochondrial matrix, cytosol and chloroplast stroma of *P. patens*. **A, C, E** Time-series show sensor responses during the first 6 h under physiological conditions and after addition of 5, 20 or 50 µM MtPQ. Measurements were performed using three-day-old protonema cultures. For each well, 360 µL protonema cultures were transferred to a 96-well microplate in KNOP ME medium. Five baseline measurement cycles were recorded before treatment addition, after which 40 µL treatment solution was added to a final volume of 400 µL per well. For the physiological control, 40 µL KNOP ME medium without stressor was added. Sensor calibration was performed using 10 mM DTT for full reduction and 10 mM $H_2O_2$ for full oxidation. Measurements were performed for **A** mitochondrial matrix-targeted sensor (MPP$_{TP}$-roGFP2-Grx1 #47), **C** cytosolic sensor (Grx1-roGFP2 #39), and **E** stroma-targeted sensor (TK$_{TP}$-Grx1-roGFP2 #40). Fluorescence was measured using a CLARIOstar plate reader with bottom optics. Ratios are shown as normalized 400-10 nm/482-16 nm excitation ratios with emission detected at 530-40 nm. Ratios were normalized to the mean of the five baseline measurements recorded before treatment addition. Blue and red horizontal lines indicate the fully reduced and fully oxidized sensor reference states, respectively. Dotted vertical lines indicate the time point of treatment addition. Markers (*) in the time-series plots indicate the time points used for the quantification. **B, D, F** Quantification of sensor ratios directly after treatment addition and after 6 h in **B** mitochondrial matrix, **D** cytosol, and **F** in chloroplast stroma. MtPQ induced a dose-dependent increase in the sensor ratio, with the strongest response observed after 50 µM MtPQ treatment. For box plots, boxes indicate the interquartile range, horizontal lines indicate medians, whiskers indicate the data range, and individual points represent replicates; n = 5 per treatment and compartment. Statistical analysis was performed separately for each compartment using two-way ANOVA to test the effects of treatment condition and time point, followed by Tukey's multiple comparisons test. Different lowercase letters indicate statistically significant differences between groups; $p < 0.05$ was considered statistically significant.

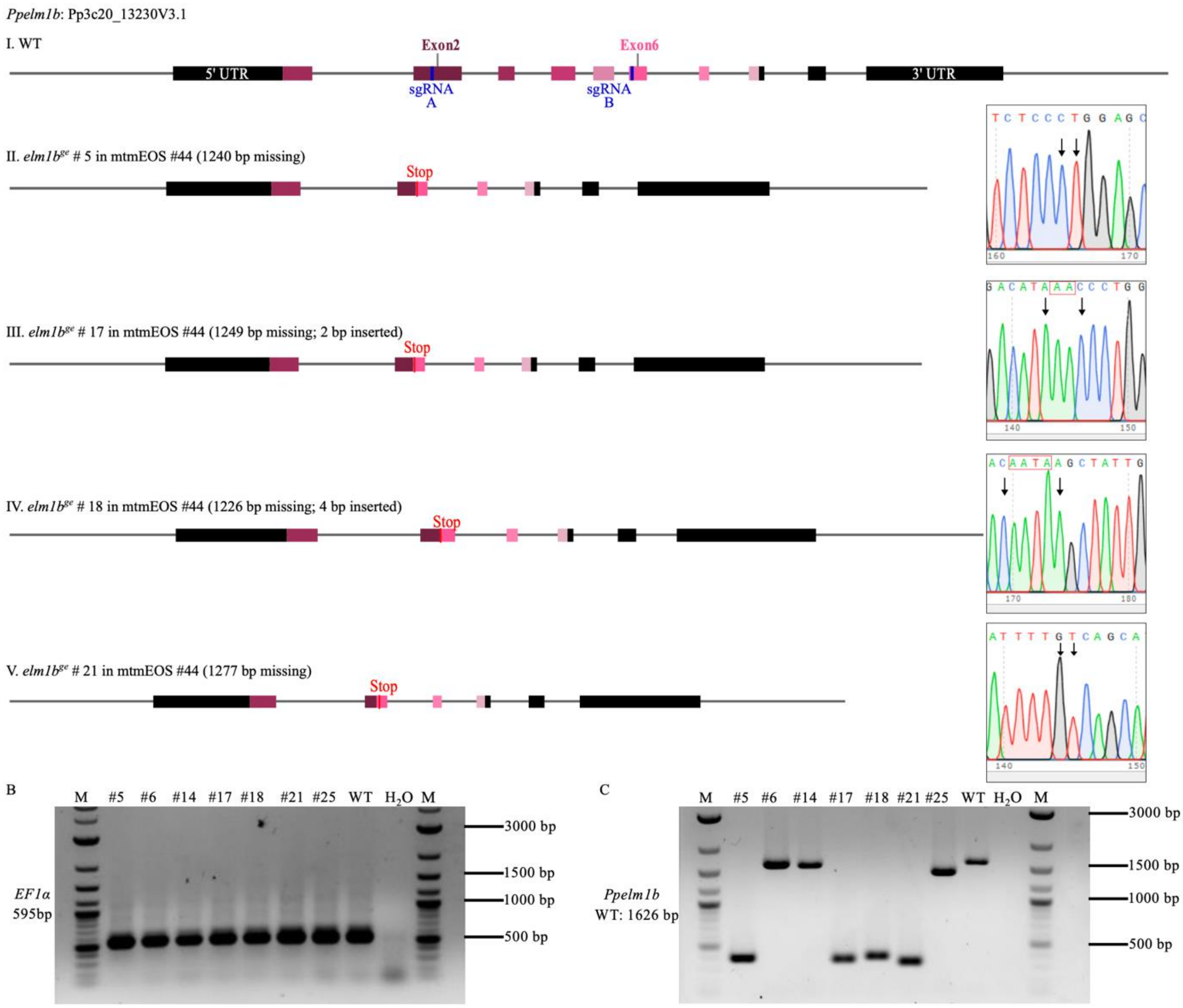


**Supplementary Figure S5: Generation and screening of CRISPR/Cas9-mediated *Ppelm1b$^{ge}$* lines in the mtmEOS #44 background.**

CRISPR/Cas9-mediated genome editing was used to generate *Ppelm1b$^{ge}$* lines in the mitochondria-targeted mtmEOS #44 reporter background. **A** Schematic representation of the *Ppelm1b* (*Pp3c20_13230*) gene structure and representative edited alleles. The wild-type locus is shown with the 5′ UTR, exons and 3′ UTR. Two sgRNAs (A and B) were designed to target the coding region. Representative edited alleles obtained in the mtmEOS #44 background are shown below the wildtype locus. The edited lines contained deletions of different sizes, in some cases accompanied by small insertions. The deletion caused frameshifts and premature stop codons, predicting truncated PpELM1B proteins. Deletions at the CRISPR/Cas9 target sites were confirmed by Sanger sequencing and are indicated by arrows. **B** PCR amplification of the reference gene Elongation Factor 1α (*EF1α*, 595 bp) was used as a control for DNA quality and template integrity. **C** PCR-based genotyping of the *Ppelm1b* target region. The wildtype allele produced a 1626 bp fragment, whereas edited lines showed shorter PCR products corresponding to deletions. $H_2O$ served as a negative control, and M indicates the DNA size marker (1 kb Plus DNA Ladder, NEB).

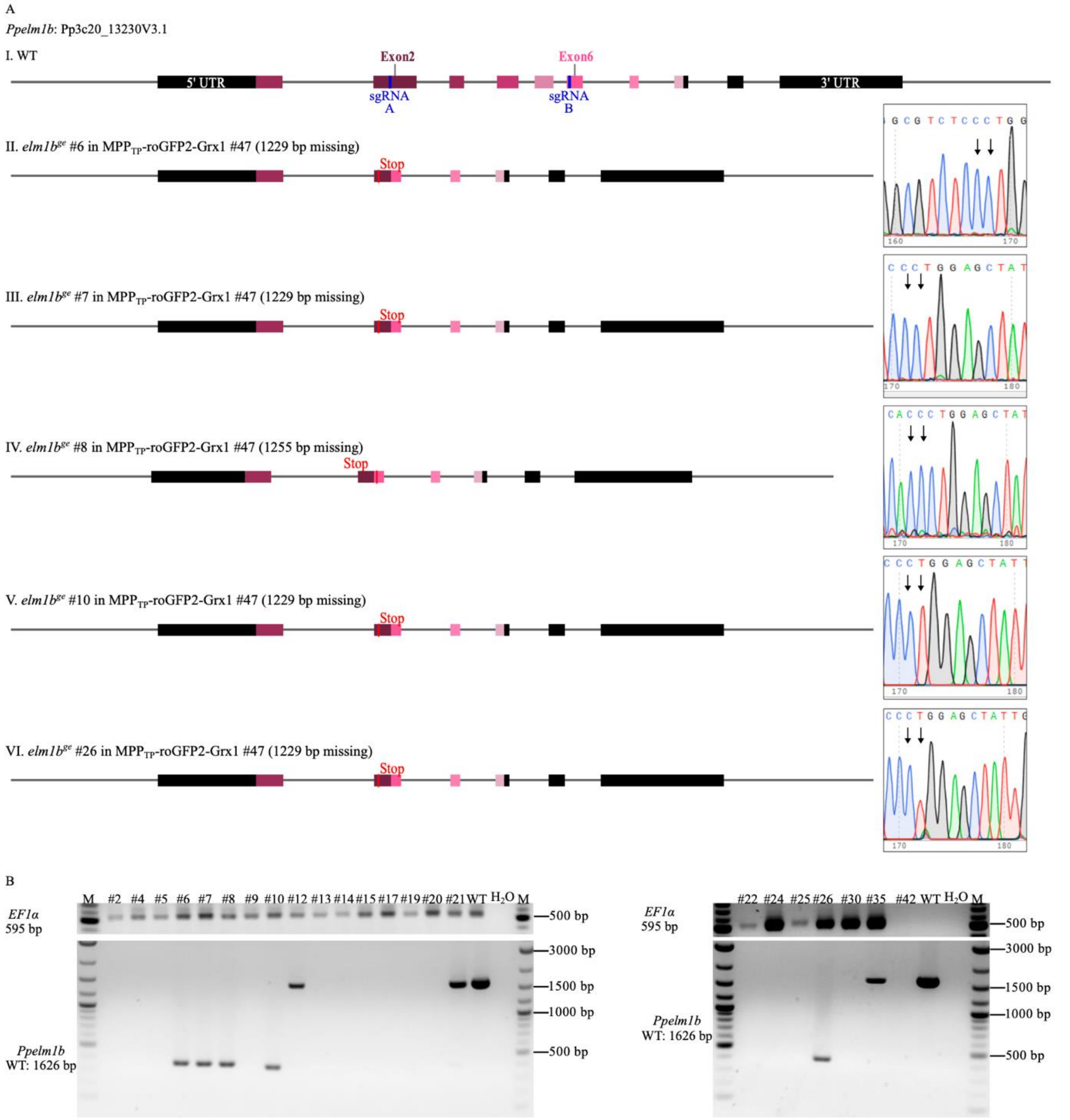


**Supplementary Figure S6: Generation and screening of CRISPR/Cas9-mediated *Ppelm1b^ge* lines in the MPP$_{TP}$-roGFP2-Grx1 #47 background.**

CRISPR/Cas9-mediated genome editing was used to generate *Ppelm1b^ge* lines in the mitochondria-targeted MPP$_{TP}$-roGFP2-Grx1 #47 reporter background. **A** Schematic representation of the *Ppelm1b* (*Pp3c20_13230*) locus and representative edited alleles generated in the MPP$_{TP}$-roGFP2-Grx1 #47 background. Two sgRNAs (A and B) were designed to target the coding region of *Ppelm1b*. Representative edited alleles are shown below the wildtype locus. The edited lines contained deletions of different sizes. All detected mutations caused frameshifts and premature stop codons, predicting truncated PpELM1B proteins. Deletions at the CRISPR/Cas9 target sites were confirmed by Sanger sequencing and are indicated by arrows. **B** PCR amplification of the reference gene Elongation Factor 1α (*EF1α*, 595 bp) was used as a control for DNA quality and template integrity. PCR-based genotyping of the *Ppelm1b* target region was performed using locus-flanking primers. The wildtype allele produced a 1626 bp fragment, whereas edited lines showed shorter PCR products corresponding to deletions in the target region. $H_2O$ served as a negative control, and M indicates the DNA size marker (1 kb Plus DNA Ladder, NEB).

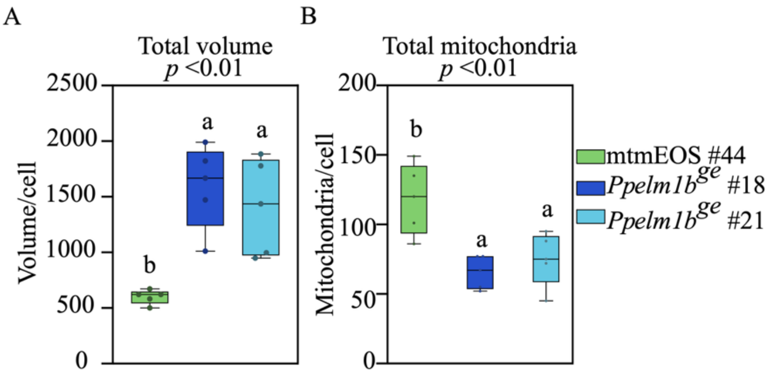


**Supplementary Figure S7: Loss of ELM1B alters total mitochondrial volume and number in *P. patens.***
Quantification of mitochondrial morphological parameters in the mtmEOS #44 background line and two independent *Ppelm1b$^{ge}$* lines (#18 and #21) by analysing confocal z-stacks in Fiji using the mitochondria analyzer plugin. **A** Total mitochondrial volume per cell. **B** Total number of mitochondria per cell. For box plots, boxes indicate the interquartile range, horizontal lines indicate medians, whiskers indicate the data range, each dot represents one cell, n = 5. Statistical analysis was performed using one-way ANOVA followed by Tukey's multiple-comparisons test. Different letters indicate statistically significant differences among groups ($p < 0.05$).

A

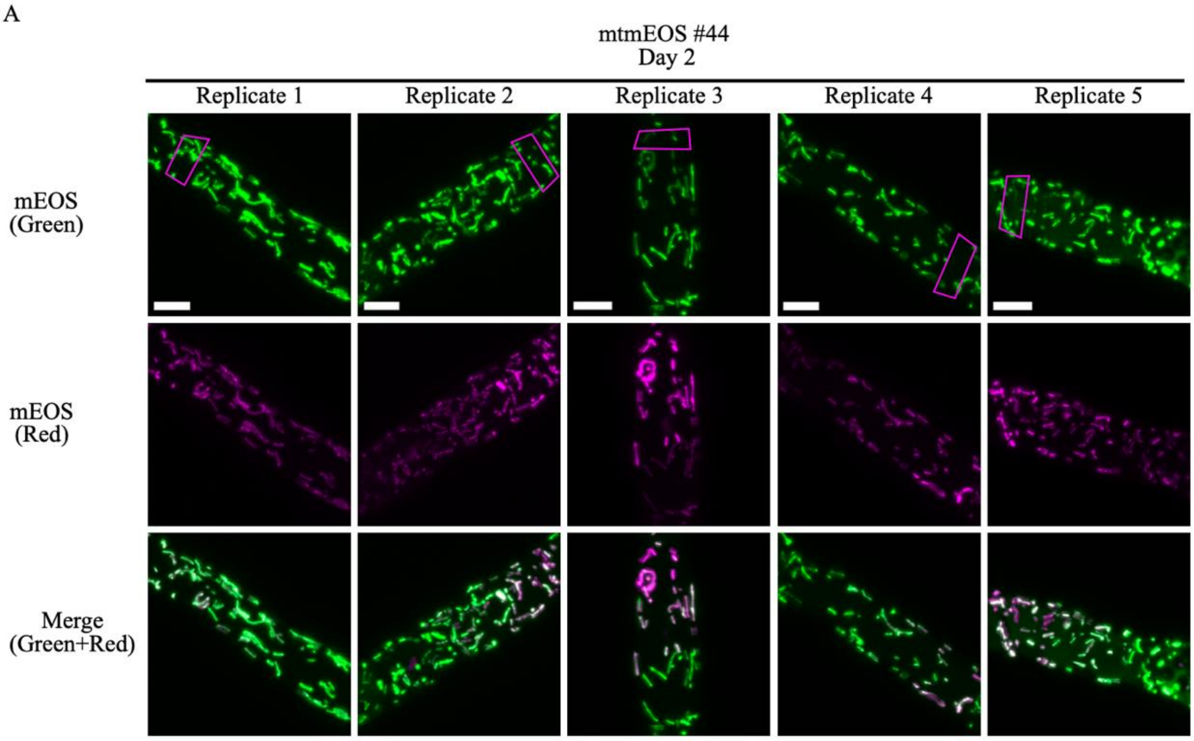


B

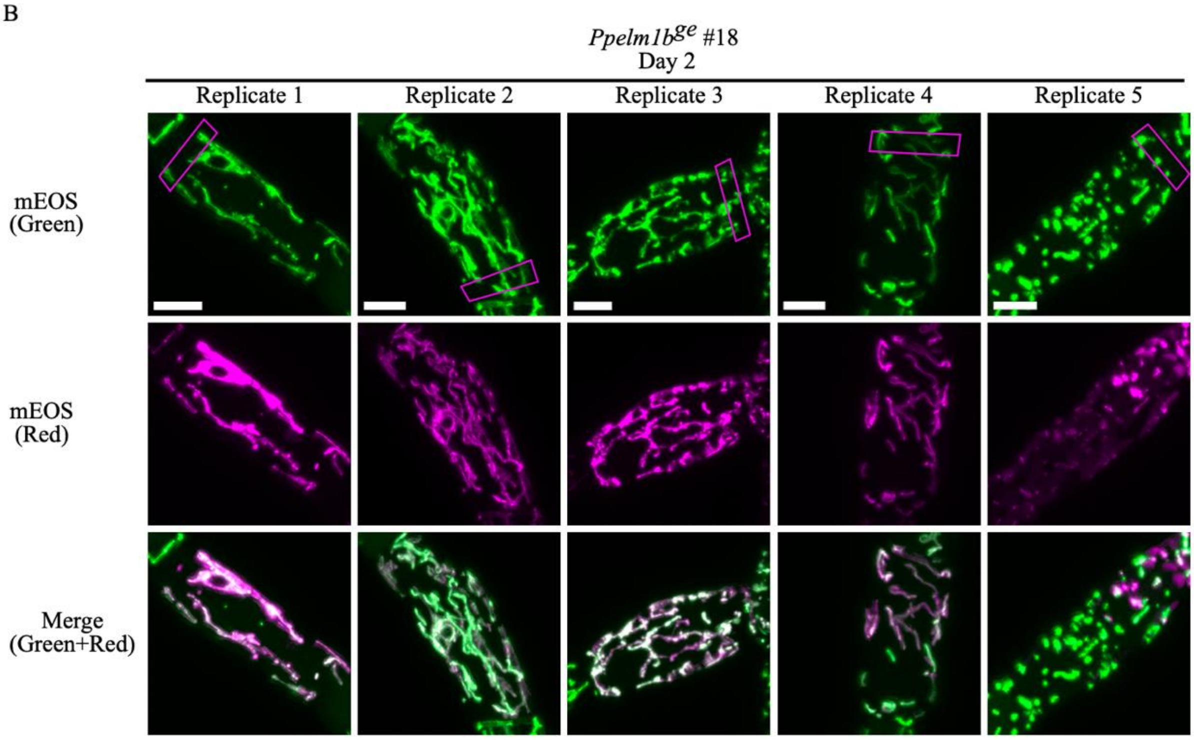


**Supplementary Figure S8: Replicate images of mEOS signal redistribution on day 2 after photoconversion.** Representative confocal images showing mitochondrial mEOS signal redistribution on day 2 after local photoconversion in the mtmEOS #44 background and in the *Ppelm1b^ge* #18 line. For repeated imaging, protonema cultures were immobilized in low-melting agar prepared in KNOP ME medium in gridded chamber slides (ibidi) and covered with fresh KNOP ME medium during imaging. Approximately 10 % of each cell was locally photoconverted on day 1, and the same immobilized cells were followed over time. Pink boxes in the $mEOS_{green}$ panels indicate the region that was photoconverted on day 1. **A** Five replicate protonema cells of the mtmEOS #44 background line imaged on day 2. **B** Five replicate protonema cells of the *Ppelm1b^ge* #18 line imaged on day 2. Maximum intensity projections (MIPs) generated from confocal z-stacks are shown. For each replicate, the green channel shows $mEOS_{green}$, the magenta channel shows photoconverted $mEOS_{red}$, and the merged image shows the overlap between both signals. In mtmEOS #44 cells, the photoconverted $mEOS_{red}$ signal spread beyond the initially photoconverted region, but the magenta channel still showed a visible gradient, with higher signal intensity near the photoconverted area. In *Ppelm1b^ge* #18 cells, the photoconverted signal appeared more evenly distributed across elongated mitochondrial structures, consistent with the quantification shown in Figure 4. Scale bars = 10 µm.

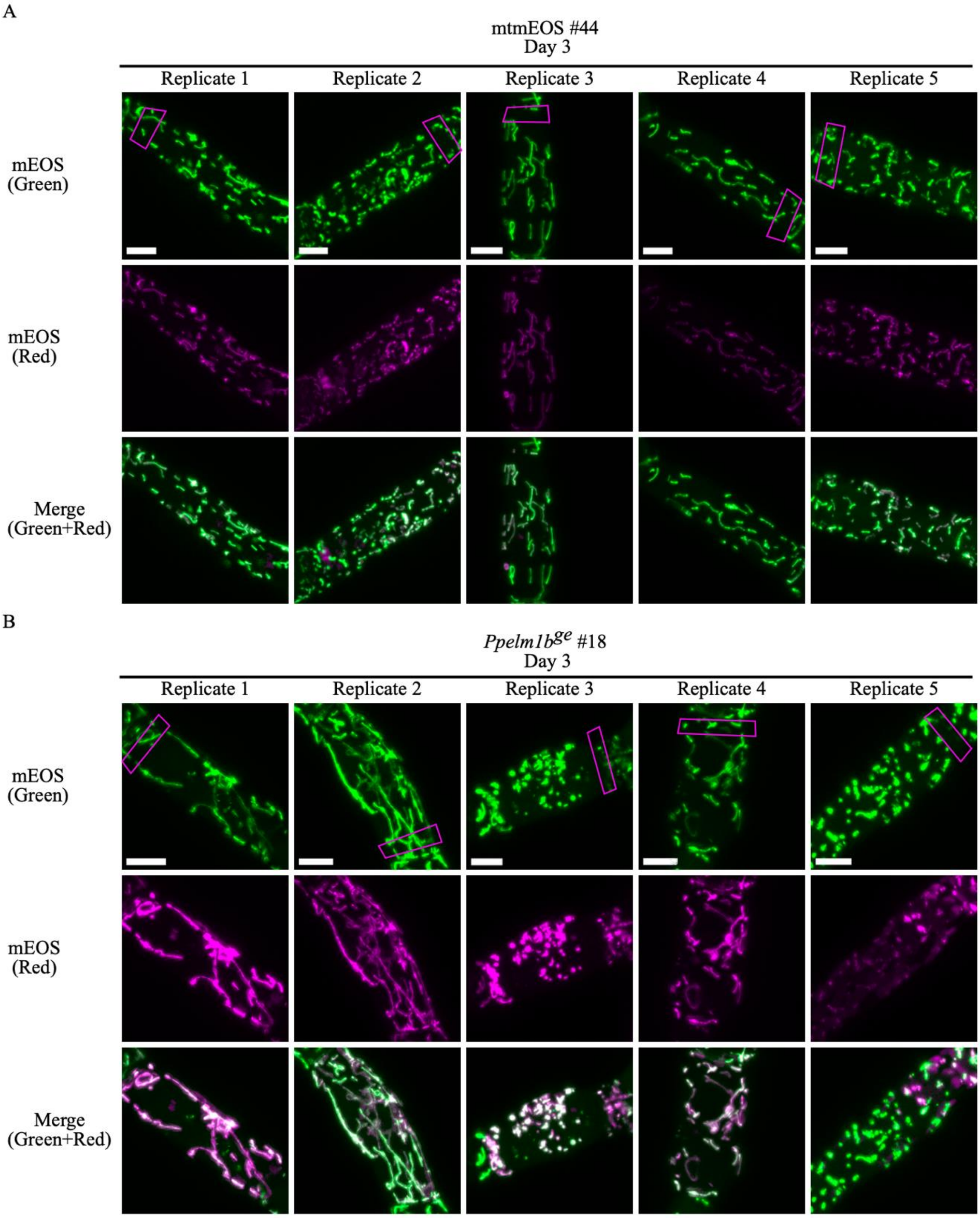


**Supplementary Figure S9: Replicate images of mEOS signal redistribution on day 3 after photoconversion.** Representative confocal images showing mitochondrial mEOS signal redistribution on day 3 after local photoconversion in the mtmEOS #44 background and in the *Ppelm1b*$^{ge}$ #18 line. For repeated imaging, protonema cultures were immobilized in low-melting agar prepared in KNOP ME medium in gridded chamber slides (ibidi) and covered with fresh KNOP ME medium during imaging. Approximately 10 % of each cell was locally photoconverted on day 1, and the same immobilized cells were followed over time. Pink boxes in the $mEOS_{green}$ panels indicate the region photoconverted on day 1. **A** Five replicate protonema cells of the mtmEOS #44 background line. **B** Five replicate protonema cells of the *Ppelm1b*$^{ge}$ #18 line. Maximum intensity projections (MIPs) generated from confocal z-stacks images. The green channel shows $mEOS_{green}$, the magenta channel shows photoconverted $mEOS_{red}$, and merged images show overlap between both signals. By day 3, the photoconverted $mEOS_{red}$ signal in mtmEOS #44 cells appeared more broadly distributed than on day 2, and the gradient from the initially photoconverted region was no longer clearly visible. In *Ppelm1b*$^{ge}$ #18 cells, the photoconverted signal remained broadly distributed across elongated mitochondrial structures, consistent with the quantification shown in Figure 4. Scale bars = 10 µm.

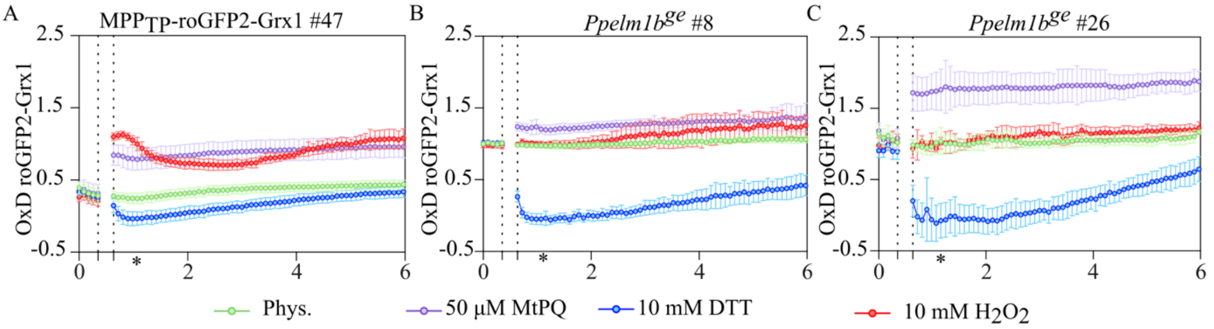


**Supplementary Figure S10: Plate reader-based comparison of *Ppelm1b$^{ge}$* lines with the mitochondria-targeted roGFP2-Grx1-expressing background.**

Plate reader-based fluorimetry was used to compare the oxidation degree (OxD) of the mitochondrial glutathione redox sensor roGFP2-Grx1 in the MPP$_{TP}$-roGFP2-Grx1 #47 background and in two independent *Ppelm1b$^{ge}$* lines generated in this background. **A-C** Time-series showing the oxidation degree (OxD) of the mitochondria-targeted roGFP2-Grx1 sensor in the MPP$_{TP}$-roGFP2-Grx1 #47 background **A** and in the independent *Ppelm1b$^{ge}$* lines #8 **B** and #26 **C**. Measurements were performed using three-day-old protonema cultures. For each well, 360 µL protonema cultures were transferred to a 96-well microplate in KNOP ME medium. Five baseline measurement cycles were recorded before treatment addition, after which 40 µL treatment solution was added to a final volume of 400 µL per well. For the physiological control, 40 µL KNOP ME medium without stressor was added. Protonema cultures were measured under physiological conditions and after treatment with 10 mM DTT, 10 mM $H_2O_2$ or 50 µM MtPQ. DTT and $H_2O_2$ were used to fully reduce and fully oxidize the sensor for OxD calculation, whereas MtPQ was used as a mitochondria-targeted oxidative stress treatment to assess the redox response of the *Ppelm1b$^{ge}$* lines. Fluorescence was measured using a CLARIOstar plate reader with bottom optics. The roGFP2 excitation ratio was calculated from fluorescence after excitation at 400-10 nm and 482-16 nm, with emission detected at 530-40 nm. OxD was calculated using the fully reduced and fully oxidized sensor states obtained after DTT and $H_2O_2$ treatment. Dotted vertical lines indicate the time point of treatment addition. Asterisks in the time-series plots indicate the time points used for quantification in Figure 5. Lines show mean values and error bars indicate standard deviation (SD), n = 5 per line and treatment.

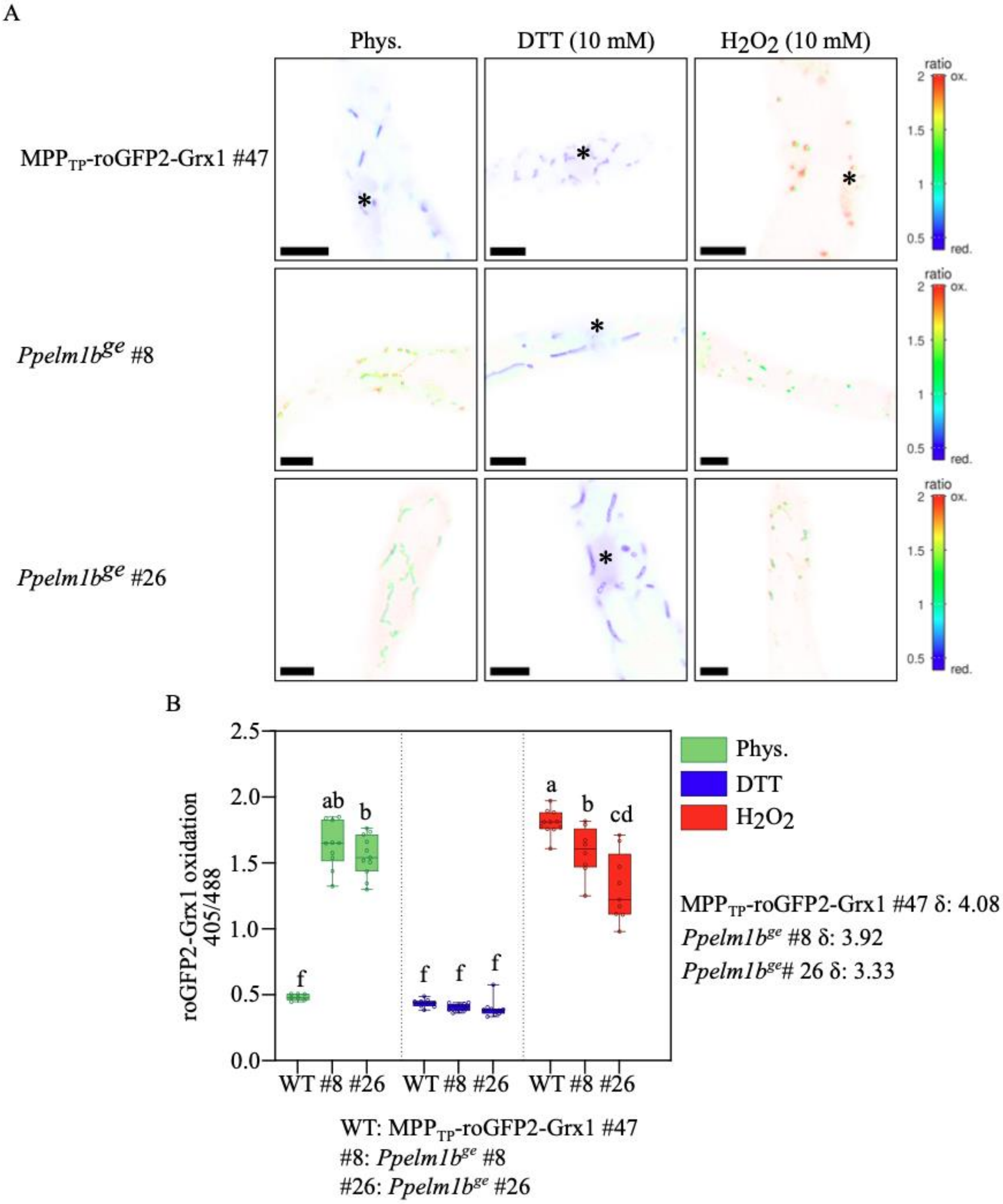


**Supplementary Figure S11: Calibration of mitochondria-targeted roGFP2-Grx1 in *P. patens*.** Calibration of the mitochondrial roGFP2-Grx1 sensor was performed in the $MPP_{TP}$-roGFP2-Grx1 #47 background and in two independent *Ppelm1b$^{ge}$* lines. Protonema cultures expressing the mitochondria-targeted sensor were imaged *in vivo* under physiological conditions and after treatment with 10 mM DTT for full reduction of the sensor or 10 mM $H_2O_2$ for full oxidation of the sensor. roGFP2 was excited sequentially at 405 nm and 488 nm, and emission was detected at 509–544 nm. **A** Representative confocal microscopy ratio images of the $MPP_{TP}$-roGFP2-Grx1 #47 background and two independent *Ppelm1b$^{ge}$* lines, #8 and #26, are shown in false-colour scale based on the 405/488 excitation ratio. Asterisks mark regions in which roGFP2-Grx1 signal was also observed in nuclei. Scale bars = 10 µm. **B** Box plots show quantification of the sensor response based on Redox Ratio Analysis (Fricker et al., 2016). Ratio values were extracted from manually defined mitochondrial regions of interest from confocal microscopy images. The measured dynamic range is indicated by δ. Boxes indicate the 25th to 75th percentiles, the centre line indicates the median, whiskers indicate the minimum and maximum values, and individual data points are shown as circles; n = 7–12. Statistical analysis was performed using two-way ANOVA followed by Tukey's multiple comparisons test. Different lowercase letters indicate statistically significant differences between groups; $p < 0.05$ was considered statistically significant.

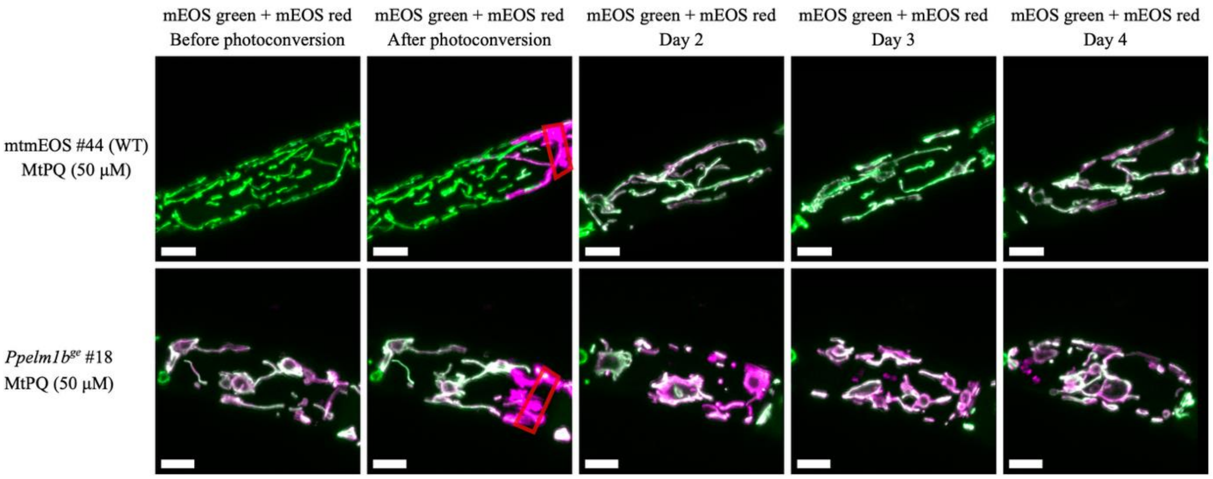


**Supplementary Figure S12: Representative confocal images of mtmEOS #44 background line and the *Ppelm1b*$^{ge}$ #18 line with 50 µM MtPQ.**
Protonema cultures were immobilized in 0.6% low-melting agar prepared in KNOP ME medium with 50 µM MtPQ and imaged repeatedly over four days. On day 1 approximately 10% of the selected cell area was photoconverted (red box), and the same immobilized cells were imaged daily. Maximum-intensity projections (MIPs) were generated from confocal z-stacks. Images are shown as merged channels, with $mEOS_{green}$ displayed in green and $mEOS_{red}$ in magenta. MtPQ treatment caused more pronounced changes in mitochondrial morphology including dognut shapes in the *Ppelm1b*$^{ge}$ #18 line from day 1 after 2 h of incubation compared with the mtmEOS #44 background. Scale bar = 10 µm.

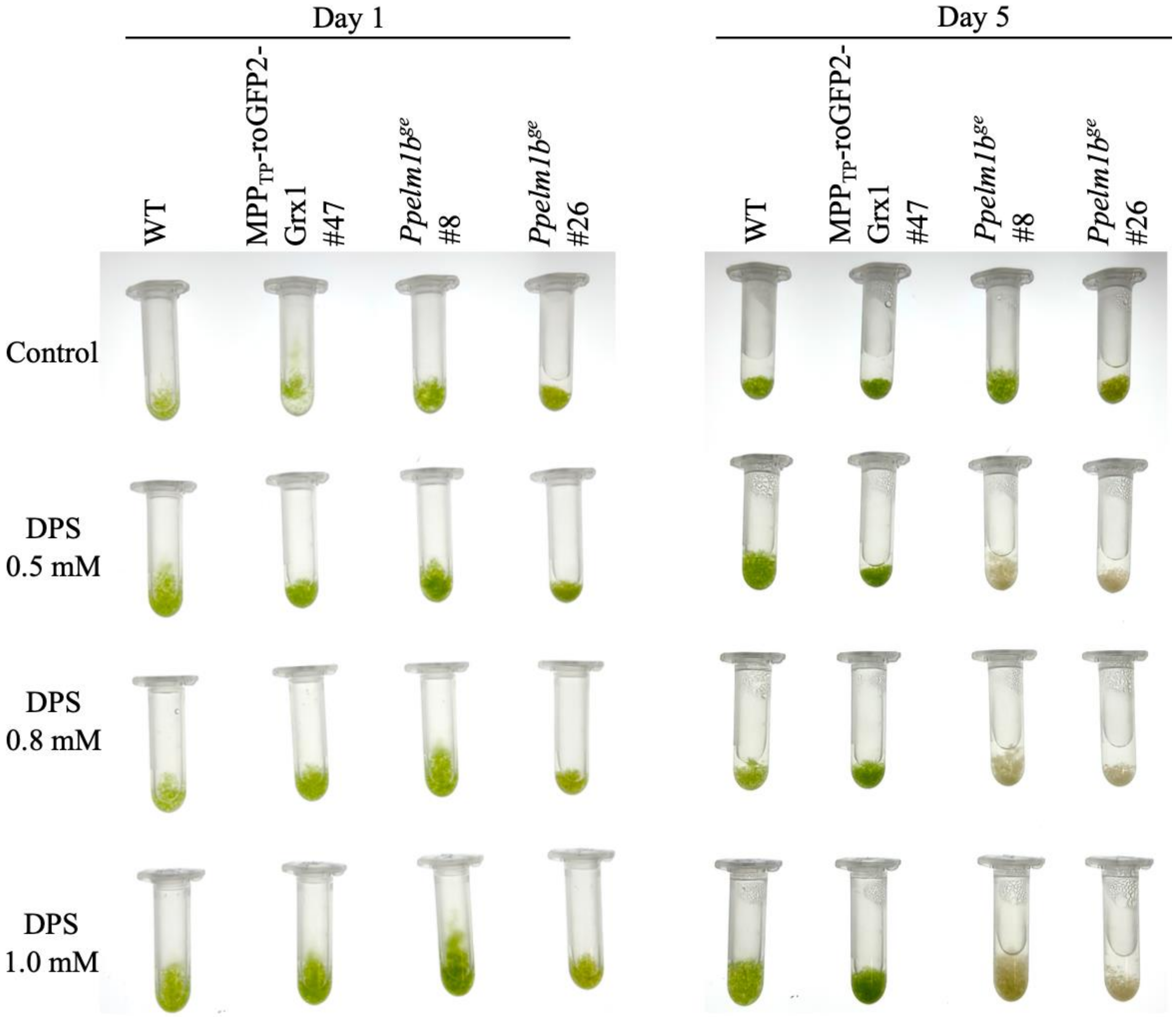


**Supplementary Figure S13: *Ppelm1b$^{ge}$* lines show increased sensitivity to DPS treatment.**
Representative images showing the response of WT, the MPP$_{TP}$-roGFP2-Grx1 #47 background line, and two independent *Ppelm1b$^{ge}$* lines, #8 and #26, to 2,2′-dipyridyl disulfide (DPS). Protonema cultures were transferred to 2 mL reaction tubes in a final volume of 1 mL KNOP ME medium containing the respective treatment. Samples were maintained under standard growth conditions and imaged daily. Cultures were kept under physiological control conditions or treated with 0.5, 0.8 or 1.0 mM DPS. Representative images are shown for day 1 and day 5 after treatment. On day 1, all cultures appeared largely green across genotypes and treatment conditions. By day 5, control cultures remained green, whereas DPS-treated *Ppelm1b$^{ge}$* #8 and *Ppelm1b$^{ge}$* #26 cultures showed visible bleaching, indicating increased sensitivity of the gene-edited lines to DPS treatment.

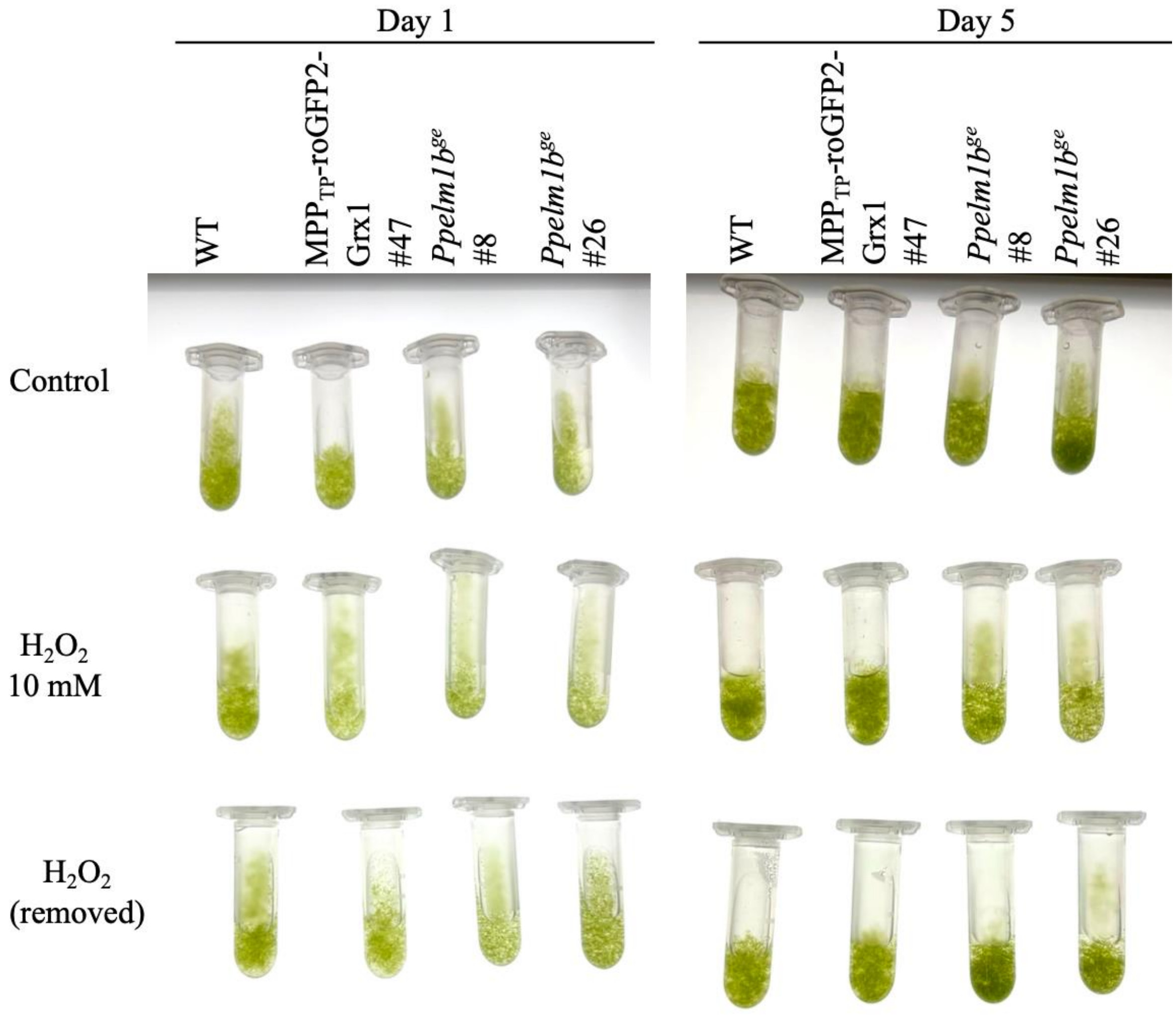


**Supplementary Figure S14: *Ppelm1b$^{ge}$* lines did not show visible sensitivity to $H_2O_2$ treatment.**
Representative images showing the response of WT, the $MPP_{TP}$-roGFP2-Grx1 #47 background line, and two independent *Ppelm1b$^{ge}$* lines, #8 and #26, to hydrogen peroxide ($H_2O_2$). $H_2O_2$ was used as a diffusible oxidative stress to test whether the sensitivity of the *Ppelm1b$^{ge}$* lines was also observed under general oxidative stress conditions distinct from MtPQ-induced mitochondrial redox cycling and DPS-induced thiol oxidation. Liquid protonema cultures were maintained under standard growth conditions and treated either continuously with 10 mM $H_2O_2$ or transiently with 10 mM $H_2O_2$ for 30 min, followed by washing and further incubation in fresh medium. Representative images are shown for day 1 and day 5 after treatment. On day 1, all cultures appeared largely green under control and $H_2O_2$-treated conditions. By day 5, control cultures remained green, and neither continuous nor transient $H_2O_2$ treatment caused any visible bleaching comparable to MtPQ or DPS treatment. Under the tested conditions, the *Ppelm1b$^{ge}$* lines therefore did not show strong visible sensitivity to $H_2O_2$.

**Table S1: Oligonucleotides**

| **Oligonucleotide** | **Sequence (5’ → 3’)** | **Target** | **Purpose** |
|---|---|---|---|
| ELM1_protosp_A_FW | CCATTAGTCGATTATTAGGCGATT | Pp3c3_340V3.1 | sgRNA A for *Ppelm1a* |
| ELM1_protosp_A_RV | AAACAATCGCCTAATAATCGACTA | Pp3c3_340V3.1 | sgRNA A for *Ppelm1a* |
| Pp_ELM1_FW_B | CCATTTATGGGGAAGACTTAGCTC | Pp3c3_340V3.1 | sgRNA B for *Ppelm1a* |
| Pp_ELM1_REV_B | AAACGAGCTAAGTCTTCCCCATAA | Pp3c3_340V3.1 | sgRNA B for *Ppelm1a* |
| Pp_ELM1S_FW_A | CCATTGGCGGCGTCTCCCACTCCC | Pp3c20_13230V3.1 | sgRNA A for *Ppelm1b* |
| Pp_ELM1S_REV_A | AAACGGGAGTGGGAGACGCCGCCA | Pp3c20_13230V3.1 | sgRNA A for *Ppelm1b* |
| Pp_ELM1S_FW_B | CCATATACAGGGAAGACTTAGCCC | Pp3c20_13230V3.1 | sgRNA B for *Ppelm1b* |
| Pp_ELM1S_REV_B | AAACGGGCTAAGTCTTCCCTGTAT | Pp3c20_13230V3.1 | sgRNA B for *Ppelm1b* |
| Pp_ELM1A_screening_fw | TTTATCTTTATTTCAGCTGTTTGAG | Pp3c3_340V3.1 | Screening of *Ppelm1a*$^{ge}$ |
| Pp_ELM1A_screening_rev | CATTTAGGTACTGATTTCGGTGT | Pp3c3_340V3.1 | Screening of *Ppelm1a*$^{ge}$ |
| Pp_ELM1B_Screening_Fw | CTCACCACAGAATGTTACGT | Pp3c20_13230V3.1 | Screening of *Ppelm1b*$^{ge}$ |
| Pp_ELM1B_Screening_Rev | CACCACAAACACGTACAATT | Pp3c20_13230V3.1 | Screening of *Ppelm1b*$^{ge}$ |
| Pp_ELM1A_short_rev | TGAAACTCACAAGCCTCTAC | Pp3c3_340V3.1 | Screening of *Ppelm1a*$^{ge}$ |
| Pp_elm1a_protoB_short_FW | GCACGTCGGTTCATTGAGTT | Pp3c3_340V3.1 | Screening of *Ppelm1a*$^{ge}$ |

**Table S2. Shape descriptors used for mitochondrial profiling**

| **Feature ID** | **Description** |
|---|---|
| Volume | Physical volume of the segmented mitochondrion, reflecting overall size. |
| Surface Area | Surface area of the segmented object, capturing boundary extent. |
| Surface area/ Volume | Surface area divided by volume, higher values indicate thinner or more convoluted shapes. |
| Major axis length | Length of the major axis of the best fitting ellipsoid, reflecting elongation. |
| Minor axis length | Length of the minor axis of the best fitting ellipsoid, reflecting thickness or width. |
| Aspect ratio | Ratio of major to minor axis length, higher values indicate more elongated mitochondria. |
| Feret diameter | Maximum caliper distance between boundary points, capturing the longest span of the object. |
| Solidity | Volume divided by convex hull volume, lower values indicate concavity or fragmentation. |
| Sphericity | Similarity to a sphere based on surface area and volume, lower values indicate deviation from spherical shape. |
| Form factor | Compactness metric derived from volume and surface area, lower values indicate less compact or more irregular shapes. |
| Extent | Fraction of the bounding box occupied by the object, describing how fully it fills its bounding box. |
| Fractal dimension | Measure of boundary or shape complexity, higher values indicate greater structural complexity. |
| Lacunarity | Measure of gap or heterogeneity in the shape, higher values indicate more uneven texture or internal void structure. |
| Skeleton length | Total length of the medial axis skeleton, approximating the length of the mitochondrial backbone. |
